# Adaptive response to BET inhibition induces therapeutic vulnerability to MCL1 inhibitors in breast cancer

**DOI:** 10.1101/711895

**Authors:** Gonghong Yan, Heping Wang, Augustin Luna, Behnaz Bozorgui, Xubin Li, Maga Sanchez, Zeynep Dereli, Nermin Kahraman, Goknur Kara, Xiaohua Chen, Yiling Lu, Ozgun Babur, Murat Cokol, Bulent Ozpolat, Chris Sander, Gordon B. Mills, Anil Korkut

**Affiliations:** Department of Bioinformatics and Computational Biology, UT MD Anderson Cancer Center, Houston, TX, 77030, USA; cBio Center, Department of Data Sciences, Dana Farber Cancer Institute, Boston, MA 02215, USA; Department of Genomic Medicine, UT MD Anderson Cancer Center, Houston, TX 77030, USA; Department of Molecular and Medical Genetics, Oregon Health and Science University, Portland, OR 97201, USA; Computational Biology Program, Oregon Health and Science University, Portland, OR 97239, USA; Department of Cell Biology, Harvard Medical School, Boston, MA 02115, USA; Department of Cell, Development and Cancer Biology, Knight Cancer Institute, Oregon Health and Sciences University, Portland, OR 97201, USA; Knight Cancer Institute, Oregon Health and Science University, Portland, OR 97201, USA; Department of Experimental Therapeutics, The University of Texas M.D. Anderson Cancer Center, Houston, TX 77030, USA; Axcella Health, Cambridge, MA 02139, USA

## Abstract

The development of effective targeted therapies for the treatment of basal-like breast cancers remains challenging. Here, we demonstrate that BET inhibition induces a multi-faceted adaptive response program leading to MCL1 protein-driven evasion of apoptosis in breast cancers. Consequently, co-targeting MCL1 and BET is highly synergistic in *in vitro* and *in vivo* breast cancer models. Drug response and genomics analyses revealed that MCL1 copy number alterations, including low-level gains, are selectively enriched in basal-like breast cancers and associated with effective BET and MCL1 co-targeting. The mechanism of adaptive response to BET inhibition involves upregulation of critical lipid metabolism enzymes including the rate-limiting enzyme stearoyl-CoA desaturase (SCD). Changes in the lipid metabolism are associated with increases in cell motility and membrane fluidity as well as transitions in cell morphology and adhesion. The structural changes in the cell membrane leads to re-localization and activation of HER2/EGFR which can be interdicted by inhibiting SCD activity. Active HER2/EGFR, in turn, induces accumulation of MCL1 protein and therapeutic vulnerability to MCL1 inhibitors. The BET protein, lipid metabolism and receptor tyrosine kinase activation cascade is observed in patient cohorts of basal-like and HER2-amplified breast cancers. The high frequency of MCL1 chromosomal amplifications (>30%) and gains (>50%) in basal-like breast cancers suggests that BET and MCL1 co-inhibition may have therapeutic utility in this aggressive subtype.

## Introduction

Despite the success of targeted therapies in cancer treatment, response to mono-therapies has been transient due the almost inevitable emergence of resistance. Blocking the routes that lead to the emergence of resistance with combination therapy is the most promising strategy to date (1-2). However, discovery of effective combination therapies is a daunting task due to complexity of drug response landscapes. A key mechanism of resistance to targeting oncogenic processes is the adaptive activation of compensatory processes, e.g. feedback loops in the short term or oncogenic alterations in the long term (3-7). Systematic approaches are needed to identify adaptive responses, reveal drug-induced vulnerabilities, and thus enable discovery of effective rational combination therapies.

Basal-like breast cancers make up nearly 20% of breast cancer cases, yet treatment options are limited for this aggressive subtype (8). This necessitates the need for the development of effective targeted therapies. The only recurrent aberrations providing therapeutic opportunities in the basal-like subtype are in the homologous recombination (HR) pathway, including BRCA1/2 (9). Gene copy number alteration of MCL1, an anti-apoptotic mediator, represents an additional potential driver aberration that is recurrent in breast cancer with an enrichment in the basal-subtype (10). With the introduction of specific inhibitors, MCL1 has become an actionable therapeutic target. Broader impact of MCL1 inhibitors could be achieved by selectively inducing survival dependencies that make use of higher levels of MCL1 (11-12). For example, targeted agents that can generate therapeutic stress and MCL1-driven anti-apoptotic dependence may induce a clinically relevant vulnerability even in cases that carry low-level MCL1 gains. Such a combination therapy strategy would be valuable from a translational perspective as it would significantly expand the number of patients that can benefit from the MCL1-based therapy while improving depth and duration of response.

Here, we demonstrate, through computational modeling and subsequent experimental validation, that BET inhibition induces an adaptive survival mechanism converging on MCL1-dependent evasion of apoptosis. The Bromodomain and Extra-Terminal (BET) proteins, including BRD2, BRD3 and BRD4, represent promising targets based on preclinical and clinical data (13-15). BET proteins link epigenetic states to lineage-specific gene expression through binding to acetylated histones and regulating transcriptional elongation downstream of enhancer sites (Figure 1A). Preclinical efficacy of BET-inhibitors (BETis) has led to clinical trials in diverse cancer types including triple-negative breast cancers (TNBC) (16) (Figure 1B). Reduction in cellular robustness, cell cycle progression, DNA damage response and induction of replication stress have been proposed as underlying mechanisms of response to BET inhibition (15, 17-18). BET molecules also regulate cellular plasticity as exemplified by the emergence of drug-tolerant persistent cells with BRD4 dependence and altered differentiation states after MEK and PI3K/mTOR inhibition in TNBC (19). The adaptive responses and therapeutic stress such as DNA-damage associated with inhibition of BET activity provide an unexplored opportunity to discover drug-induced vulnerabilities, which might be exploited with effective combination therapies (20).

**Figure 1.**
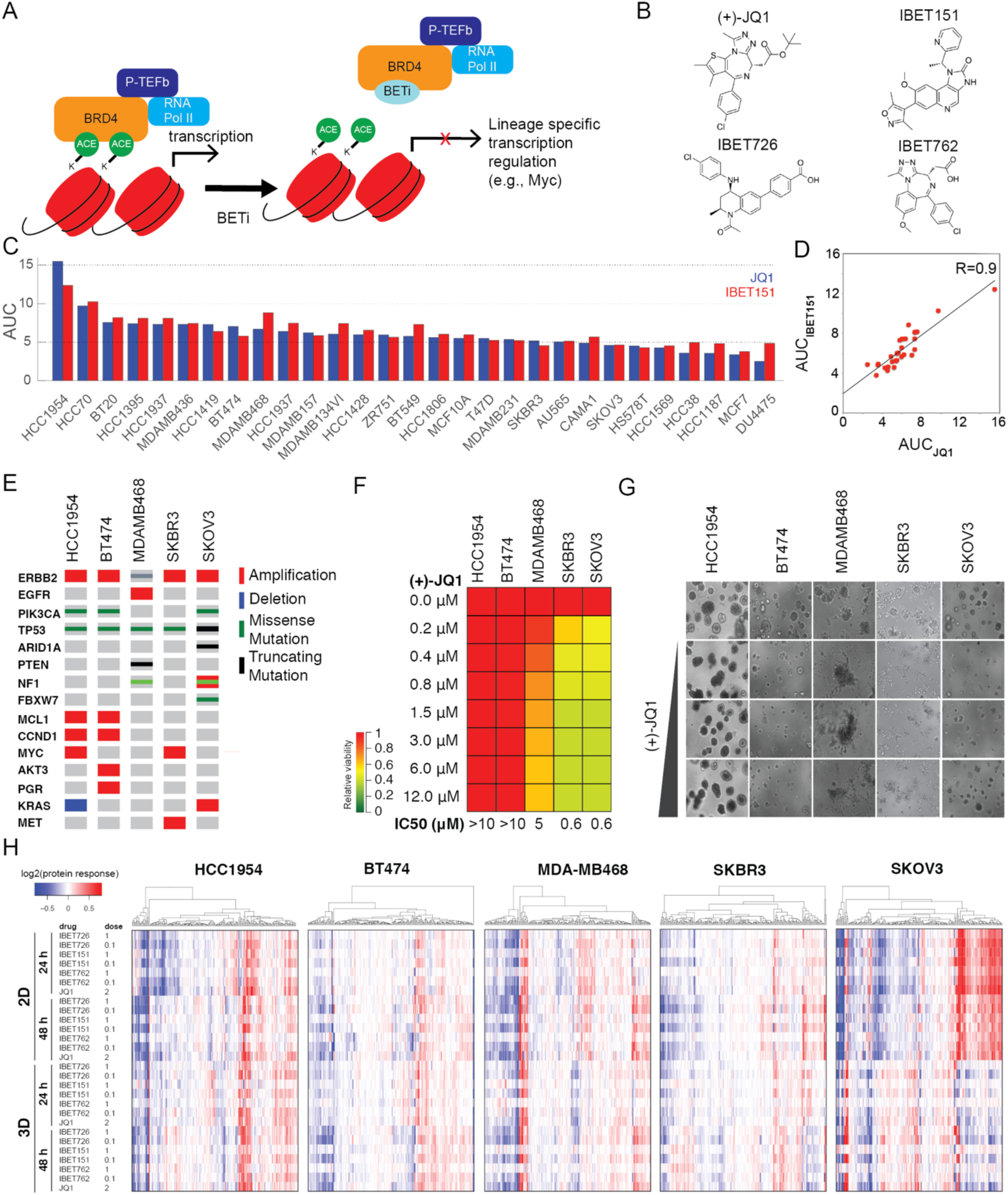
Response to BET inhibition in breast cancer. **A.** BETis bind to the acetyl histone binding cavity on BRD4 and prevent recruitment of BRD4 to chromatin and block expression of lineage specific genes including MYC. **B.** Chemical structures of BETis used in perturbation experiments. **C.** The relative responses to the BETis JQ1 and IBET151 across 28 breast cancer lines and the ovarian cancer line SKOV3 are quantified as area under the curve (AUC) of the dose-response relationship (dose range 0-10 **μ**M). **D.** Scatter plot of responses to IBET151 vs. JQ1 across the cell lines in (C) demonstrates similarity in phenotypic response to the two BETis. **E.** Landscape of potential driver oncogenic events in selected breast and ovarian cancer cell lines. **F.** Dose-dependent responses to JQ1 in cell lines cultured in 3D matrigel medium. **G.** Images of dose-dependent responses to JQ1 in 3D matrigel cultured cells. **H**. Proteomic response map of cell lines to BET inhibition. Proteomic data was collected with RPPA using antibodies that quantify 217 total or phosphoprotein levels (two time points, 2D and 3D cultures, varying doses of the four BETis, colors represent the degree of proteomic responses to inhibition in log2 space).

We demonstrate that combined targeting of MCL1 and BET proteins is synergistic in multiple in vivo and in vitro breast cancer models. We identified a mechanism of a BETi induced adaptive response program that involves activation of lipid metabolism coupled to increased membrane fluidity, cellular motility and receptor tyrosine kinase signaling. The adaptive response program leads to increased levels of the anti-apoptotic protein MCL1 and subsequent evasion of apoptosis, which confers resistance to BET inhibition, and yet, vulnerability to MCL1 inhibition. Genomics and cell viability analyses revealed that chromosomal amplifications or low-level copy number gains in the MCL1 gene are predictive of sensitivity to the combination of MCL1 and BET inhibitors. This combination may have therapeutic value as MCL1 copy number alterations, either as high-level amplifications or low-level gains, are frequent in breast cancer. Our integrated approach provides a generalizable protocol for identification of effective precision combination therapies in diverse cancer types.

## Results

### Breast cancer cells have differential responses to BET inhibition

To address the efficacy spectrum, predictors of response, and mechanisms of resistance to BET inhibition, we profiled responses to BET targeting in 28 breast and one ovarian cancer cell line. The effects of BET inhibition on cell viability (Figure S1A) were quantified by the area under the curve (AUC) of dose-response relations (Figure 1C) and IC50 (Figure S1B) for JQ1 and IBET151, two structurally different and efficacious BET inhibitors. A comparison of responses to JQ1 and IBET151 revealed a high correlation (R=0.91), suggesting that both compounds have similar efficacies and on-target activity *in vitro* (Figure 1D). Consistent with previous studies, a correlative genomic analysis of response to BET inhibition did not detect a significant enrichment for any recurrent/driver breast cancer genomic aberration within resistant or sensitive cell lines (Figure S1A) (8,21-22). The lack of genomic markers predictive of response to BETi in breast cancer necessitates deeper mechanistic analyses to discover biomarkers likely to identify patients who will benefit from BETi.

We tested cell viability responses to BETi in both 2D monolayers and 3D spheroid cultures in 5 selected cell lines. The cell lines (ordered from most resistant to sensitive), HCC1954 (HER2 amplified, basal-like), BT474 (HER2+/ER+), MDAMB468 (basal-like, HER2-/EGFR amplified), SKBR3 (HER2+) and SKOV3 (HER2+, ovarian), were selected to cover different levels in the BETi sensitivity spectrum. Despite being an ovarian cancer line, SKOV3 was selected as a sensitive control line to decode events exclusive to more resistant cells for the following reasons: First, SKOV3 originates from a patient with serous adenocarcinoma, an ovarian cancer subtype with similarities to basal-like breast cancers (8). Second, SKOV3 also carries major genomic similarities to breast cancers samples (e.g., TP53-null, HER2 amplified, PIK3CA-mutated). The comparison of responses in 2D vs. 3D media determines whether matrix attachment alters responses to BET inhibition (Figure 1E-G). The impact of 3D culture on cell viability was most dramatic for the most sensitive cell line, SKOV3, in which IC50 increased from ∼200nM to 1.5μM (Figure S1C).

We interrogated proteomic response patterns to identify processes that could potentially explain the observed sensitivities to BETis. Using reverse phase protein arrays (RPPA), we measured changes in 217 key oncogenic signaling molecules in response to four different BETis (JQ1, IBET151, IBET726, and IBET762) at two time points (24 and 48 hours) in 2D monolayer or 3D matrix-attached spheroid cultures of the five cell lines (Figure 1H). The RPPA targets were chosen to monitor activity and levels of representative molecules from key pathways including PI3K, RAS-MAPK, Src/FAK, RTK signaling axes, DNA repair, cell cycle, apoptosis, immuno-oncology, and histone modifications (23). The phosphoproteomic response to BET inhibition in diverse cell lines, time points, and conditions generate a comprehensive response map of the BETi-based epigenetic perturbations in breast cancer cells (Figure S1D-E).

### Computational network modeling identifies MCL1 upregulation as an adaptive response to BET inhibition

We analyzed the BETi proteomic response map using our Target Score network modeling method (Figure 1, see methods). Our goal is to identify collective adaptive responses that can drive drug resistance and induce drug-induced vulnerabilities to combination therapies (Figure 2A-B). Based on a reference network model of signaling (Figure 2C, S2A), we calculated the target scores for all proteins after treatment with JQ1, IBET151, IBET726, and IBET762 compounds in HCC1954, BT474, MDMB468, SKBR3, and SKOV3 lines using phosphoproteomic data collected 24 and 48 hours after drug perturbation.

**Figure 2.**
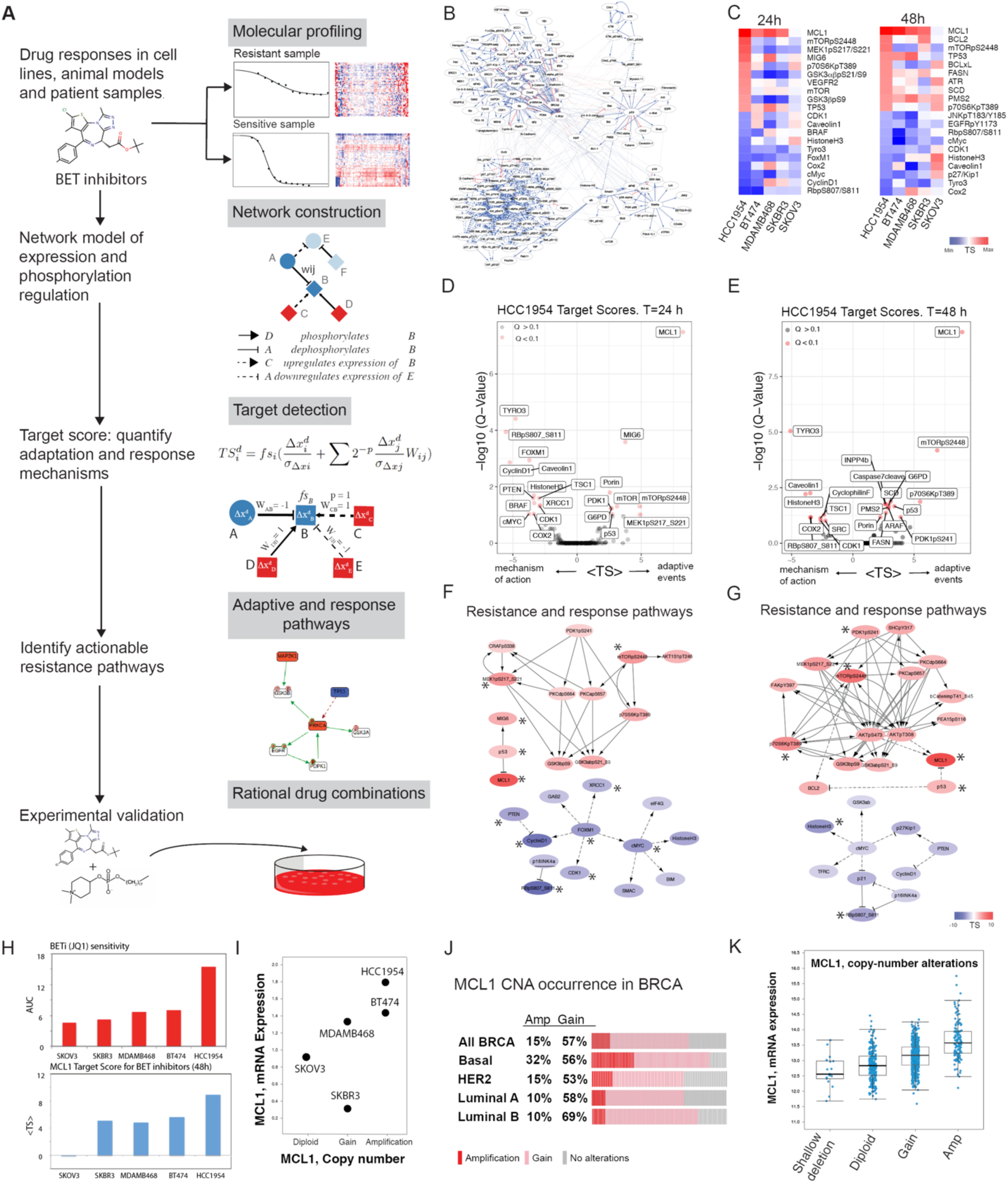
The computational Target Score analyses of adaptive responses to BET inhibition in breast cancer. **A**. The Target Score algorithm (see methods for details) identifies network-level adaptive responses to targeted perturbations. The method involves, 1. Molecular profiling of responses to perturbations in samples with varying sensitivity to the perturbation agent. 2. Construction of a reference network that captures potential relations between all measured proteomic entities. 3. Quantification of a sample-specific adaptation (target) score that links protein interactions to drug response on the reference network using proteomic drug response data. 4. Identification of network modules that have a significantly high target score (i.e., collectively participate in adaptive responses) in each sample. 5. Selection of actionable targets that participate in adaptive responses and experimental testing of drug combinations. **B.** Reference network model constructed for 217 phosphoproteomic entities (see methods for construction of the model, Figure S2 for the network diagram in high resolution). **C.** Network-level adaptive responses are quantified as target scores using the reference network and BETi response data. Heatmaps capture the scores for 24 (left) and 48 (right) hours post-treatment in 3D cultures. The proteomic entities with the highest and lowest 10 target scores in HCC1954 are used as the reference and cell lines are ordered from most resistant (HCC1954) to most sensitive (SKOV3). **D, E.** Analysis and statistical assessment of adaptive responses (high target score) and direct responses (low target score) to JQ1, IBET151, IBET726, and IBET762 in HCC1954 24 hours and 48 hours post-treatment. Target scores are the average values across the four BETis. The Q-values represent the FDR-adjusted P-values against the target score null distribution (see methods). **F.** Network modules of adaptive responses (red nodes representing high target score) and mechanisms of action (blue nodes representing low target score) 24 hours post-treatment. * denotes the significantly strong target scores according to the statistical validation (See methods) **G.** Network modules of adaptive responses and mechanisms of action for BET inhibition, 48 hours post-treatment. **H.** Association of MCL1 target score (top) and BETi sensitivity (bottom). The target score bar chart represents the mean target score across four BETis. **I.** copy number and mRNA expression status of MCL1 in cell lines. **J.** The frequency of breast cancer samples with high (red) and low (pink) level copy number amplifications in MCL1. **K.** Distribution of MCL1 mRNA levels with varying MCL1 copy number status across breast cancer samples (TCGA).

Differential analysis of target scores for JQ1 across cell lines identified adaptive responses that are either exclusive to most resistant cell types or shared across all samples (Figure 2C, S2B). In the most resistant line, HCC1954, the anti-apoptotic protein MCL1 had the highest target score value in response to JQ1, in both 24 and 48 hours among the 217 proteins and phosphoproteins assessed. In HCC1954, 24 hours after JQ1 treatment, Target Score identified an additional series of molecules downstream of receptor tyrosine kinase (RTK) signaling such as mTOR, mTOR_pS2448, MIG6, MEK1/2_pS217/S221, P70S6K_pT389, GSK3**α**/**β** _pS21_S9, and AKT1S1_pT246. The enrichment suggests involvement of RTK signaling in the adaptive response. At 48 hours, more diverse adaptive responses were observed as proteins involved in apoptosis (MCL1, BCL2, BCL-XL), DNA repair (ATR, PMS2), lipid metabolism (SCD, FASN), AKT/mTOR signaling (mTOR_pS2448, INPP4B) and TP53, which is mutated in HCC1954, had high target scores. Target scores in BT474 carried important similarities to HCC1954 (Figure S2B). In BT474, MCL1 had a high target score (5^th^ highest at 24 hours and the highest at 48 hours). At both 24- and 48-hour time points, high target scores in BT474 were associated with total protein level changes in EGFR and its downstream targets (Figure S2C). In conclusion, Target Score analysis nominated the upregulation of anti-apoptotic MCL1 molecule as a key adaptive response to BET inhibition exclusively in resistant lines likely as a consequence of altered EGFR signaling and lipid metabolism.

To detect a statistically robust and consistent adaptive response signature, we analyzed the average target scores for each protein across four different BETis (Figures 2D, 2F). We eliminated high target scores that are solely driven by network connectivity biases without significant input from cell type-specific response data using a bootstrapping based statistical assessment method. The method generates a null model of target scores using randomized data sets (mixed protein labels) and the reference network model. On the null model, we identify the target scores with high values and low FDR-adjusted P-value (Q<0.1), an indication that the scores are driven by both the network and sample specific data (see methods). MCL1 (at both 24hr and 48hr), MIG6 (at 24hr), and mTORpS2448 (at 48hr) had the most significant (Padj < 10^−4^) and highest target scores in HCC1954. At both time points, multiple markers of AKT pathway activation (P70S6K_pT389, PDK1_pS241, mTOR_PS2448), MAPK (MEK1/2_pS217/S221), and lipid metabolism (SCD and FASN) pathways had high target scores. The FDR-adjusted p-values decreased to less than 0.1 at 48 hours particularly for the lipid metabolism mediators SCD and FASN. MCL1 target score followed a near monotonously decreasing trend with increasing sensitivity to JQ1 assuming the lowest target score in SKOV3 (most sensitive) line and highest target score in HCC1954 (most resistant) (Figure 2H, Figure S2). The trend suggests MCL1 upregulation is correlated with resistance to BET inhibition. Although mTOR_pS2448 also had high target scores in HCC1954, there was no correlation between the target scores and sensitivity to JQ1 across additional cell lines. In conclusion, statistical analysis results for JQ1, IBET151, IBET726, and IBET762 were consistent with JQ1-focused analyses particularly in the nomination of MCL1 as a key driver of adaptive responses (Figure S2).

In both HCC1954 and BT474, cell cycle proteins RB1_pS807, CyclinD1, c-Myc, and p27/Kip1 had the lowest target scores (Figure 2E-G, Figure S2B-D). Interestingly, in BT474, phosphorylation of MAPK and AKT pathway members were also marked with low target scores despite increases in total proteins levels and corresponding high target scores of total protein expression in EGFR, AKT and MAPK pathway members, suggesting the existence of BRD4 inhibitors altering signaling processes downstream of EGFR/HER2 in BT474. Consistent with existing literature, our differential and statistical analyses of target scores suggest that BETis reduce cell cycle progression (24-25) as indicated by low target scores in cell cycle pathways.

### MCL1 mRNA and protein levels inversely correlate with BETi sensitivity

The analysis of MCL1 copy number, transcriptomic and proteomic status provides orthogonal evidence on the roles of MCL1 in drug-resistant cells. MCL1 is copy-number-amplified and mRNA overexpressed in the resistant lines HCC1954 and BT474 (Figure 2I) (CCLE data source, 26). MCL1 is copy-number-gained and has intermediate mRNA expression in the MDAMB468 line, which has a slightly lower JQ1 resistance. More sensitive cell lines, SKBR3 and SKOV3, have low mRNA expression and/or diploid copy number states for MCL1. The MCL1 aberration pattern in cell lines reflects the distribution of MCL1 status in patient cohorts. High-level copy number amplifications and low-level gains in the MCL1 locus are observed in 15% and 57% of breast cancer cases in the TCGA cohort (N=1084 patients), respectively (Figure 2J). MCL1 aberrations are particularly enriched in the aggressive basal subtype with 32% of cases being copy number amplified. Despite the well-established role of MCL1 amplification as a breast cancer driver, little is known of the significance of low-level gains, which lead to relatively high mRNA expression compared to MCL1 diploid cases (Figure 2K).

We profiled MCL1 levels in BET-inhibitor-treated and drug-naive cells with varying JQ1 sensitivity (Figure 3A). As expected from copy number and transcriptomic states, HCC1954 and BT474 cells (MCL-1 amplified) expressed substantial levels of MCL1 protein in their drug-naive state, whereas SKOV3 cells (MCL1 wild-type) had lower MCL1 protein expression. In BT474 and HCC1954, treatment with JQ1 led to a time-dependent increase of MCL1 levels (Figure 3A). In contrast, MCL1 protein levels did not increase in BETi sensitive SKOV3 cells in response to short term (48 hours) BETi treatment. Motivated by these evidence lines, we next experimentally tested the combined effect of BET and MCL1 inhibition in breast cancer cells. We focused on lines that are most resistant to BET inhibition where predicted effects are most likely to be manifest.

**Figure 3.**
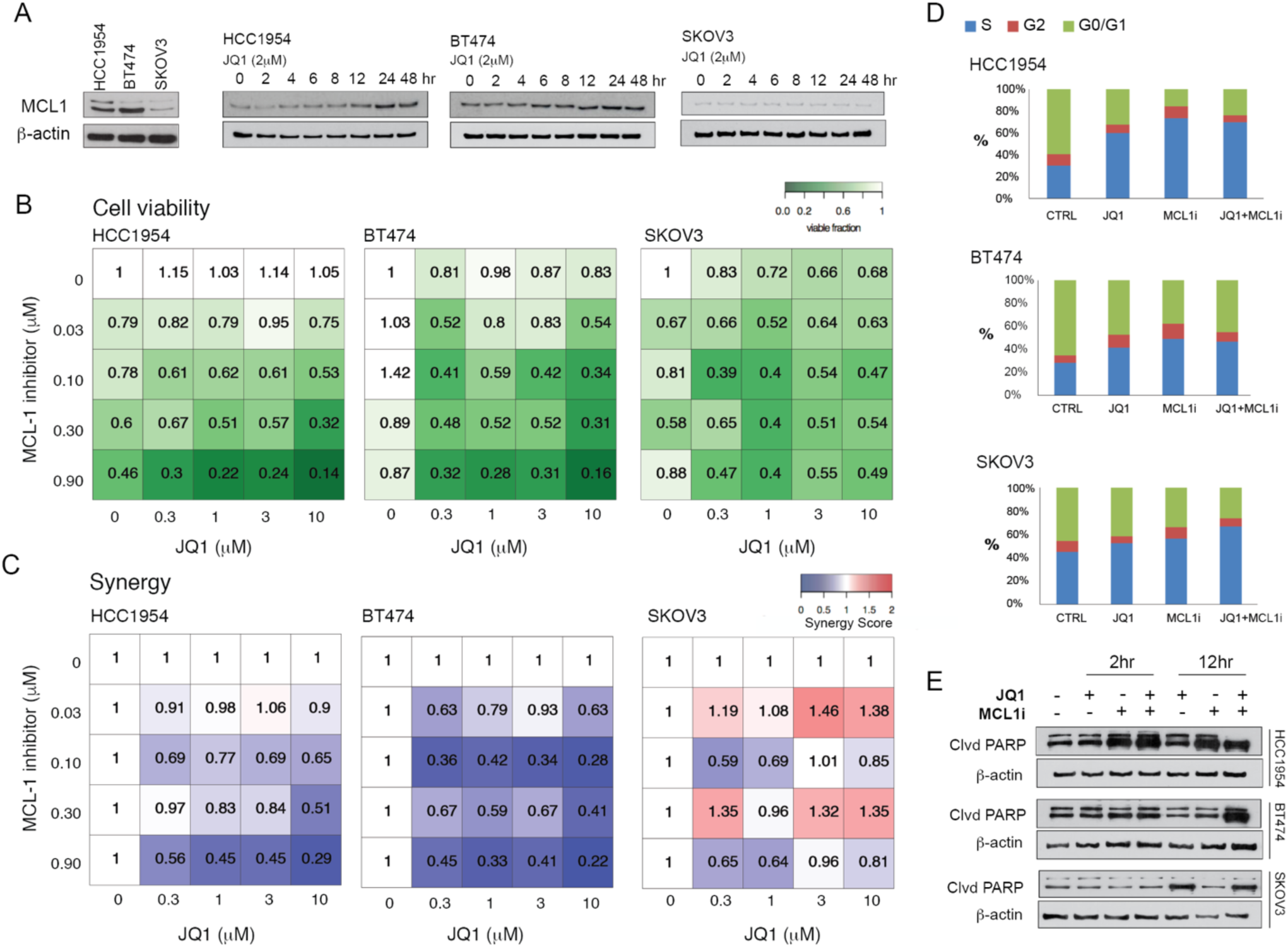
Synergistic responses to the combination of BET and MCL1 inhibitors in breast cancer. **A**. MCL1 protein expression in drug-naive BETi resistant (HCC1954, BT474) and BETi sensitive (SKOV3) cells (left). MCL1 protein level changes in response to BET inhibition (2μM of JQ1, 0 to 48 hours). **B.** Cell viability response to inhibitors of MCL1 (S63845) and BET in HCC1954, BT474 and SKOV3 cells in 3D spherical cultures supplemented with 2% matrigel. **C.** Interactions between JQ1 and S63845 are quantified using the Bliss independence method. **D**. The shift in cell cycle stage distribution in HCC1954, BT474 and SKOV3 in response to JQ1 (2μM) and S63845 (0.3μM) combination 48 hours post-treatment are quantified using flow cytometry. **E**. Western blotting analysis of cleaved PARP levels monitors apoptotic response to JQ1 and S63845, 2 and 12 hours post-treatment.

### BET and MCL1 inhibitors are synergistic in breast cancer cells

We treated BETi-resistant HCC1954 and BT474, and BETi-sensitive SKOV3 cells with combinations of JQ1 and a highly selective MCL1 inhibitor (MCL1i) S63845 (11) (Figure 3). We measured cell viability in response to the combination of MCL1 inhibitor (S63845) and BETi (JQ1) (Figure 3B). Consistent with our previous experiments, increasing doses of JQ1 in HCC1954 and BT474 did not affect cell viability. We observed a substantial response to MCL1 inhibition (0.9 μM S83645) in HCC1954, with a 54% reduction of cell viability, while BT474 was relatively more resistant with a 13% reduction. Both cell lines were highly responsive to combination treatment as evidenced by more than 80% reduction in cell viability when JQ1 and S83645 were introduced at 10 and 0.9 μM, respectively. In contrast, SKOV3 was not responsive to the S83645 and the combination did not alter the already high activity of JQ1. We quantified synergy between BETi and MCL1i using the Bliss independence metric, where lower independence scores suggest synergy and higher values suggest antagonism (Figure 3C). A Bliss score of less than 0.5 represents a strong synergy. In both BT474 and HCC1954, the combination was highly synergistic at diverse doses. In SKOV3, no synergy was observed and the two compounds were antagonistic in multiple-dose configurations with Bliss scores as high as 1.45. Thus, we validated the Target Score based predictions that co-targeting MCL1 and BET would be effective in BETi-resistant cells.

To identify mechanisms underlying synergy, we measured the effects of BET and MCL1 targeting on cell cycle progression and apoptosis. Cell cycle progression and arrest were quantified with DNA content measurements 48 hours after drug perturbations using flow cytometry (Figure 3E). Consistent with the Target Score calculations, which suggested a reduction of cell cycle pathway activities, JQ1 induced an S-phase cell cycle arrest in all samples. In HCC1954, the S-phase population enriched from 30% to 59.8% upon 2μM JQ1 treatment. In BT474, we observed a similar increase in the fraction of cells in S-phase (from 28% to 41.5%) upon JQ1 treatment. The shift, which was induced by JQ1 in the S-phase population was lowest in SKOV3, with an increase from 44.8% to 52.4% after 48 hours. Next, we monitored apoptotic responses through detection of PARP cleavage (Figure 3F). Importantly, 12 hours of JQ1 treatment at 2μM induced PARP cleavage in SKOV3 but not in drug-resistant HCC1954 and BT474. The observed cell cycle arrest across all cell lines and lack of apoptotic cells in drug-resistant cell populations explain the static cell viability profiles in response to monotherapy in HCC1954 and BT474. Single agent treatment with S83645 at 0.3μM induced PARP cleavage in HCC1954 but not in SKOV3 and BT474, which is consistent with BT474 and SKOV3 cells being refractory to single-agent MCL1 inhibition. The combination of JQ1 (BETi) and S8364 (MCLi), induced PARP cleavage in all tested cell lines, particularly in HCC1954 and BT474. The apoptotic shift and reduced cell count in response to JQ1 and S83645 combination provides strong evidence that resistance to BETi is driven by the upregulation of MCL1 activity in breast cancer (Figure 3E-F). This finding also further validates the utility of predictions from the Target Score algorithm.

### MCL1 copy number status is a predictor of response to co-targeting MCL1 and BET

Next, we investigated predictors of response to MCL1 inhibition alone or in combination with BET inhibition. We focused on MCL1 copy number status motivated by the computational predictions and the subsequent experimental validation of the drug combination in MCL1 amplified cell lines. To investigate the role of MCL1 amplification, we tested the effect of MCL1 and BET inhibitors on cell viability in 10 cell lines (9 breast lines and SKOV3) with varying cancer subtypes and genomic backgrounds (Figure 4A). The response to therapy was quantified and evaluated by (i) normalized area under the curve (nAUC = AUC_observed_/AUC_100% viability at all doses_) for drug efficacy across multiple doses, (ii) A_max_ for responses at high doses, and (iii) drug synergy (Bliss independence) for drug interactions. The weakest responses to MCL1 inhibition were observed in HCC1937, HCC1419 and SKOV3, which have either diploid or single-copy loss of the MCL1 gene (Figure 4B-C). Normalized AUC (**μ** = 0.93) and Amax (**μ** = 83%) were significantly higher in HCC1937, HCC1419 and SKOV3 compared to other lines (p-val=0.007, H_o_: nAUC^MCL1i^(MCL1_wt/hetloss_) = nAUC^MCL1i^(MCL1_gain/amp_); p-val<0.001, H_o_: A_max_^MCL1i^ (MCL1_wt/hetloss_)=A_max_^MCL1i^(MCL1_gain/amp_)). Cell lines with either low-level gain or high-level amplification of the MCL1 gene were more responsive to MCL1 inhibition with effective eradication of tumor cells (A_max_ **μ**= 0.19) at high doses and a significantly lower AUC (**μ**= 0.59). The combination of MCL1 and BETis decreased cell viability in all tested cell lines albeit with modest effects in intrinsically JQ1 sensitive lines such as HCC1937, SKOV3 and MCF7. The mean nAUC across cell lines with MCL1 copy number alterations was reduced from 0.59 for MCL1 mono-therapy to 0.33 for the combination. For cell lines with wild type or heterozygous loss of MCL1, the nAUC reduced from 0.90 to 0.60 with combination therapy. A more dramatic response was observed in A_max_ compared to AUC. In cells with MCL1 amplification, a near-complete eradication of tumor (A_max_ **μ**= 0.05) was observed in contrast to cell lines carrying no MCL1 amplification (A_max_ **μ**= 0.36) (Pval < 0.001, H_o_: A_max_^MCL1i+JQ1^ (MCL1_wt/hetloss_) = A_max_^MCL1i+JQ^(MCL1_gain/amp_)). Next, we analyzed the association between MCL1 copy number status, mRNA levels, and A_max_ for the response to JQ1 and S63845. The copy number and mRNA expression values for each cell line were extracted from the CCLE database. The analysis demonstrates MCL1-low (mRNA and copy number) cell lines are clearly separated in their responses to the combination from MCL1-high cell lines, which have much lower A_max_ (i.e., higher sensitivity) (Figure 4D). We did not observe any significant enrichment of increased MCL1i and/or MCL1i + BETi sensitivity in tumors with other recurrent oncogenic alterations (Figure S4). The separation based on MCL1 copy number and mRNA levels suggests a clear association with MCL1 levels and response to the drug combination.

**Figure 4.**
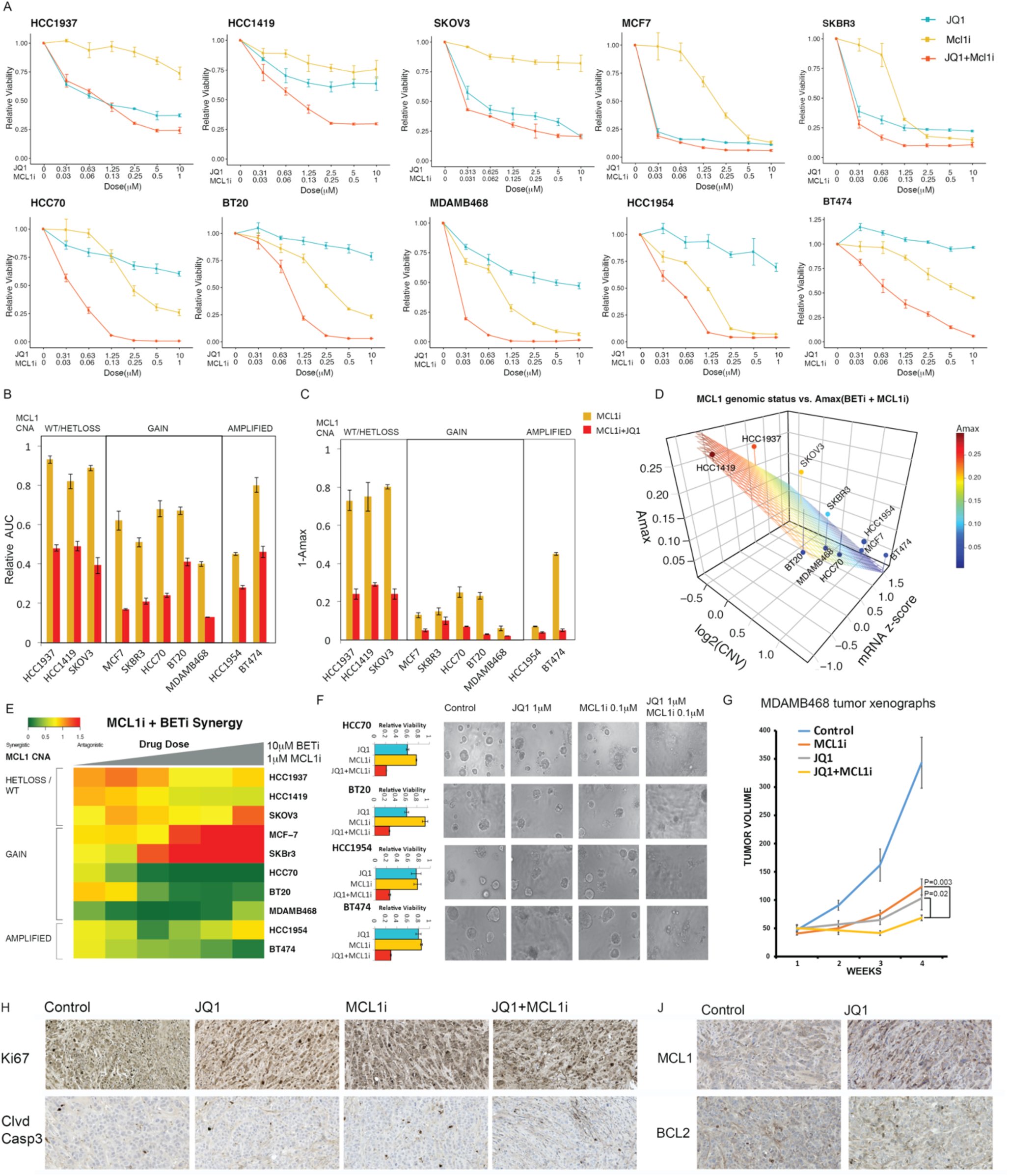
Molecular determinants of responses to MCL1 and BET inhibition. **A.** Dose-response curves of BETi (JQ1) and MCL1i (S63845) in breast and ovarian cancer cell lines. **B**. The normalized AUC (AUC/AUC_100% viability_) for S63845 and combination of JQ1 and S63845. Cell lines are classified according to MCL1 copy number status. **C.** A_max_ (response at maximum dose) for S63845(1μM) and combination of JQ1 (10μM) and S63845 (1μM), classified according to MCL1 copy number status. **D.** The relation between combination therapy A_max_, MCL1 mRNA levels and MCL1 copy number status as profiled in the CLLE project for all tested cell lines. The surface represents the linear regression between A_max_ (dependent) and MCL1 copy-number and mRNA (independent) variables across cell lines. **E.** The synergy, additivity and antagonism between JQ1 and S63845 are quantified with Bliss independence score for increasing doses of drugs for cell lines with varying MCL1 copy number status. **F.** Cell viability responses to JQ1 (1μM) and S63845 (0.1 μM) in 3D cultures of select cell lines treated with JQ1 (1μM) and S63845 (0.1 μM). Images are collected 72 hours after drug treatment. **G**. Tumor volume curves in MCL1-high MDAMB468 xenografts treated with BETi and MCL1i, each agent at 20mg/kg, i.p injections 3 times/week). N= 10/arm for control, 9/arm for treatment cohort. **H**. Representative images of IHC analysis for Ki67 and cleaved Caspase 3 in xenografts 10 days after onset of drug treatments. **J.** Representative images of IHC analysis for MCL1 and BCL2 in response to BET inhibition 10 days after onset of in vivo drug treatments (see figure S4 for MCL1 and BCL2 IHC analysis in response to MCL1i and BETi+MCL1i).

We computed drug synergy using Bliss independence score for all drug doses across all cell lines and detected three clusters with distinct BETi and MCL1i synergy (Figure 4E). The first cluster (HCC1937, SKOV3, HCC1419) consisted of MCL1 inhibitor-resistant lines and is characterized by additive drug interactions between BET and MCL1 inhibitors (0.5 < Bliss score < 1.0). In the second cluster (SKBR3, MCF7), an antagonistic interaction (Bliss score > 1.0), which can be attributed to the high efficacy of JQ1 as a single agent, was observed. In the third cluster (HCC70, BT20, MDAMB468, HCC1954, and BT474) a strong synergy was observed with Bliss synergy scores below 0.5. The cells in the third cluster were resistant to JQ1 and also carry MCL1 copy number gains. Finally, we confirmed that synergistic interactions between JQ1 and S63845 were also preserved in 3D spheroid cultures (Figure 4F). In summary, a robust and strong synergy exists between targeting MCL1 and BET in the BETi resistant and MCL1 amplified context. Thus, based on the drug responses and synergies across cell lines with varying MCL1 levels, MCL1 copy number status is a predictor of response to the drug combination that could potentially be translated to the clinic to identify patients likely to benefit.

### BET and MCL1 co-targeting is highly effective *in vivo*

On the basis of the synergy between MCL1 and BET inhibitors *in vitro*, we investigated the efficacy of the combination in the MDAMB468 xenograft model in nude athymic NCr mice. The MDAMB648 model represents the basal breast cancer subtype with an amplification in EGFR, loss of PTEN, TP53 point mutation and a low-level chromosomal gain in MCL1. Mice were treated with MCL1 and BET inhibitors as single agents and in combination at 20mg/kg intraperitoneal injections 3 times a week (n = 9 in experiment groups, n = 10 in control arm). Although both single agents generated a partial response, the combination therapy induced a significantly stronger effect with a marked decrease in tumor volume compared to each single agent after 3 weeks of treatment (two-tailed Mann–Whitney U test; p=0.003, Ho: <ΔTV^MCL1i^>= <ΔTV^MCL1i+BETi^>; p=0.017, Ho: <ΔTV^BETi^>= <ΔTV^MCL1i+BETi^>) (Figure 4G, S4B). Next, we monitored the *in vivo* changes in protein levels during the progression of therapy (day 10 after therapy onset) in key response markers with immunohistochemistry. The changes in cell proliferation and apoptotic markers *in vivo* were consistent with the *in vitro* results. The proliferation marker Ki67 were lowest in tumors of mice treated with the drug combination. Similarly, levels of the apoptosis marker, cleaved caspase 3 were highest in combination-treated samples. Cleaved caspase 3 levels were intermediate in MCL1i treated animals and lowest in BETi treated and control cases. As predicted by the computational models and in vitro experiments, MCL1 but not BCL2 protein levels were increased in response to BET targeting (Figure 4H-J, S4C). BRD4, the main target of the BET inhibitor, was ubiquitously expressed in control and drug-treated states (Figure S4D). No animal weight loss was observed in any of the treatment arms suggesting the agents had no observable toxicity as single agents or in combination (Figure S4F). In summary, BET inhibitor increased MCL1 protein levels and induced a therapeutic vulnerability to MCL1 inhibition *in vivo*.

### BETi induced MCL1 upregulation is mediated by EGFR/HER2 signaling activity

To elucidate signaling mechanisms underlying adaptive MCL1 protein upregulation, we searched for phosphorylation cascades that could link BET inhibition to MCL1 protein accumulation and subsequent apoptotic evasion. We quantified differential phosphorylation of each protein and focused on most deviant proteins based on RPPA data from resistant and sensitive lines (Figure 5A). The most differentially phosphorylated proteins were predominantly driven by the difference between the most resistant line HCC1954 and the sensitive line SKOV3. Statistical analysis revealed enrichment of HER2/EGFR downstream signaling in BETi resistant cells as phosphorylation of HER2, EGFR, SHC, SRC, AKT, GSK3**α**/**β**, MEK1/2 and 4EBP1 increased in response to BET inhibition in HCC1954 but not in SKOV3. In addition to RTK signaling, we observed increased phosphorylation of Myosin II and NDRG1 in HCC1954. The differential analysis is consistent with the Target Score analyses, which identified a module of BETi adaptive response centered around MCL1 and enriched with signaling molecules downstream of EGFR/HER2 (e.g., MAPK, AKT, SHC, mTOR, PKC) (Figure 2H). Based on the results, we hypothesized that BETi induced MCL1 increase and subsequent apoptosis evasion occurred through activation of HER2/EGFR and downstream signaling.

**Figure 5.**
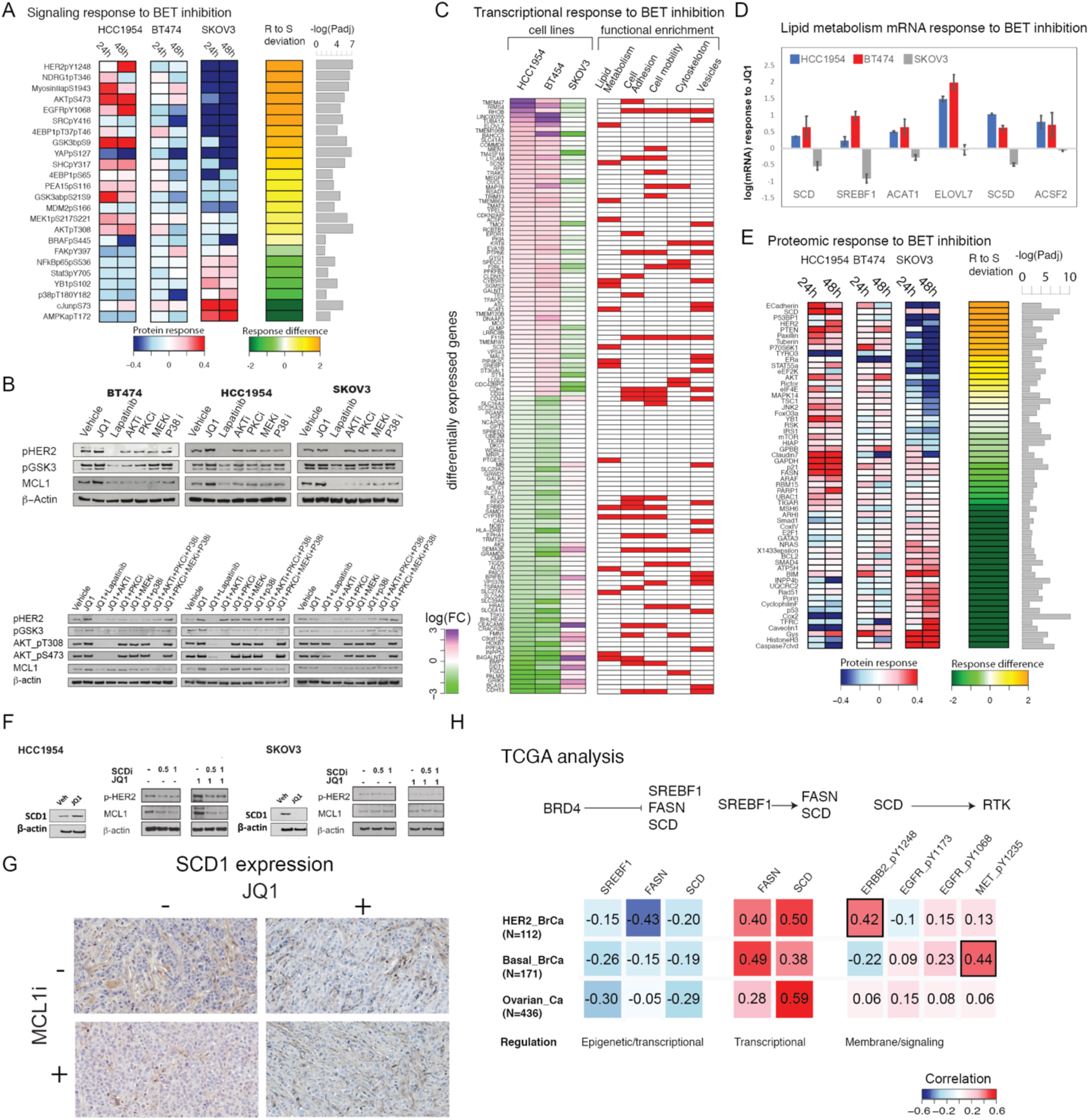
Molecular mechanisms of response to combined BET and MCL1 targeting. **A.** Differential analysis of phosphoproteomic response to BETis in resistant (HCC1954, BT474) and sensitive (SKOV3) cells. The heatmap represents the average log fold difference (log[X_perturbed_/X_unperturbed_]) across four BETis (JQ1, IBET151, IBET726 and IBET762). “R to S deviation” is the difference of fold changes between resistant (HCC1954 and BT474) and sensitive (SKOV3) cells (see methods for RPPA analysis). The proteins with significantly different expressions between HCC1954 (most resistant) and SKOV3 (sensitive line) are listed (Padj <0.05). **B.** MCL1, AKT signaling and HER2 activity changes in response to pathway and BET inhibition in HCC1954, BT474 and SKOV3 are measured. Responses to pathway inhibitors applied as single agents (top panel) and combinations with JQ1 are monitored to demonstrate the role of signaling pathways linking BET and MCL1 activity. **C.** Differential analysis of transcriptomic responses to BET inhibition in resistant vs. sensitive cell lines. The first three columns in the heatmap demonstrate fold changes of mRNA species that change in opposite directions in resistant versus sensitive lines. The red/white columns on right represent the involvement of each differentially expressed gene in different molecular processes according to a GO-term enrichment analysis. **D.** Transcriptomic responses in key genes of the lipid metabolism pathway as identified by the differential response analysis. **E**. Differential total protein analysis is performed as in (A) with a focus on only total protein changes. The heatmaps represent the proteomic responses averaged across four BETis. **F**. The analysis of SCD, MCL1 and p-EGFR/p-HER2 levels in response to inhibitors of BET (JQ1 at 1μM) and SCD (A939572). The SCD, MCL1 and p-EGFR/p-HER2 levels are measured 48 hours after drug perturbation in HCC1954 and SKOV3 cells. **G.** Representative images of IHC analysis of in vivo SCD expression changes in the mouse xenograft model (MDA-MB468) treated with BET and MCL1 inhibitors. **H.** Correlation analysis of proteomic and transcriptomic levels involved in the proposed associations linking BET to EGFR/HER2 within HER2+, basal and ovarian cancer patients. Each analysis reflects a particular step in the pathway -i.e., BET to fatty acid pathway genes, fatty acid enzyme expression, fatty acid enzymes to RTK signaling. All correlations are in the range of −0.5 to 0.5, consistent with relatively low correlations observed between omic entities in other TCGA studies. As a reference, the average correlation between mRNA and protein levels of a given gene is estimated within a range of 0.3-0.6 across the whole genome in different studies (27-28).

To test this hypothesis, we profiled MCL1 levels and signaling under systematic perturbations (Figure 5B). We treated HCC1954, BT474, and SKOV3 cells with inhibitors of BET (JQ1), EGFR/HER2 (Lapatinib), AKT (MK-2206), P38/MAPK14 (Doramapimod), PKC (Dequalinium Chloride), and MEK1/2 (trametinib) applied as single agents. In BT474 and HCC1954, both MCL1 and GSK3**α**/**β**_pS9/S21 levels increased in response to JQ1, confirming the role of MCL1 in the adaptive response, and substantially decreased in response to Lapatinib, suggesting a causal link from HER2/ERGFR to MCL1. Inhibitors of downstream effector molecules (AKT, MAPK14, MEK1/2, PKC) had either partial or no impact on MCL1 levels, particularly in HCC1954, suggesting no single path downstream of HER2/EGFR can single-handedly regulate MCL1. To test whether the inhibition of individual signaling routes downstream of EGFR/HER2 can reverse adaptive responses to BET inhibition, we treated the cells with combinations of pathway and BET inhibitors. In BT474 and HCC1954; JQ1 driven increases in MCL1, AKT_pS373/pT308 and GSK3**α**/**β**_pS9/S21 were reversed by Lapatinib consistent with the role of HER2/EGFR activation in mediating adaptive responses to BET inhibition. In contrast, MCL1 and GSK3**α**/**β**_pS9/S21 levels modestly increased (HCC1954) or did not change (BT474) in response to combinations of JQ1 with all other agents (i.e., inhibitors of AKT, MEK1/2, PKC and MAPK14/P38) suggesting that multiple signaling axes are required for the BETi adaptive response or alternatively, inhibition of one mediator results in activation of the other pathways. Indeed, adaptive responses were blocked only with a cocktail of three signaling inhibitors with JQ1 (i.e., JQ1 with cocktails of PKCi+P38i+AKTi or PKCi+P38i+MEK1/2i). The molecular responses to combinatorial perturbations suggest that EGFR/HER2 activation is necessary and sufficient for increased MCL1 in HCC1954 and BT474. However, none of the tested inhibitors other than Lapatinib could block the adaptive response to BET inhibition, indicating highly redundant downstream signaling cascades. This suggests parallel and redundant routes downstream of EGFR/HER2 converge to regulate MCL1 levels.

### A BETi induced transcriptional program alters fatty acid metabolism to activate EGFR/HER2 signaling

To explore the mechanisms of EGFR/HER2 activation by BET inhibition, we analyzed transcription and total-protein changes in response to BET inhibition. Transcriptomic responses were profiled with 1μm JQ1 treatment for 48h in HCC1954, BT474 and SKOV3 cells with mRNA sequencing (Figure 5C). The mRNA analysis identified 127 differentially responding transcripts in HCC1954/BT474 versus SKOV3 (see methods). A GO-term analysis of differentially expressed genes identified 85 statistically significant pathways with enrichment of biosynthetic pathways particularly lipid metabolism, cell adhesion, and motility in resistant lines (Supplementary Table 1). The enrichment profiles suggest that the BETi transcriptional program is associated with lipid bilayer cell membrane changes in drug-resistant samples. Stearoyl-CoA desaturase (SCD), which catalyzes the rate-limiting step in monounsaturated fatty acid (MUFA) synthesis and regulates lipid membrane fluidity in cells (29-31), was in 43 out of 85 significant pathways. In addition to SCD, other key lipid metabolism enzymes, including SREBF1 - a master transcription factor that regulates expression of key fatty acid enzymes (e.g., SCD, FASN), ELOVL7, SC5D, ACSF2, and ACAT1 are differentially overexpressed in response to BET inhibition in resistant cells (Figure 5C-D). Moreover, differential analysis of total protein expression demonstrated the involvement of cell adhesion and fatty acid metabolism in BETi response, with E-cadherin and SCD as the highest-ranked differentially expressed proteins in resistant cells (Figure 5E). BETi also increased MyosinII_pS1943 and decreased AMPKa_pT172 supporting that BET inhibition alters motility and lipid metabolism.

Motivated by the roles of SCD as a rate-limiting step in MUFA formation and a regulator of membrane fluidity, we focused on SCD involvement in modulating EGFR/HER2 activity and MCL1 protein levels in response to BET inhibition (Figure 5F). First, we validated that BET targeting leads to increased SCD protein levels in BETi resistant HCC1954 cells but not in sensitive SKOV3 cells. Treatment of HCC1954 cells with the SCD inhibitor (SCDi, A939572) decreased both p-HER2/EGFR and MCL1 levels, suggesting a role of SCD upstream of the HER2/EGFR - MCL1 signaling axis. To identify whether the BETi-induced MCL1 accumulation depends on SCD, we treated HCC1954 with JQ1 and A939572. Both HER2 phosphorylation and MCL1 decreased upon combined BET and SCD inhibition, suggesting that SCD inhibition can reverse adaptive responses to BET inhibition. Consistent with the *in vitro* models, in MDAMB468 tumor xenograft models BET inhibition but not MCL1 inhibition leads to increased SCD IHC staining relative to the untreated control condition (Figure 5G). Thus, BETi induced expression of lipid metabolism genes, especially SCD, is an intermediate event between BET inhibition and HER2/EGFR to MCL1 signaling axis both in vitro and in vivo.

Next, we explored the interplay between BET inhibition, fatty acid synthesis and RTK activation leading to MCL1 upregulation, in patient cohorts. We performed a correlation analysis of mRNA or protein levels in therapy-naive basal-like breast cancer, HER2+ breast cancer, and serous ovarian cancer TCGA data (Figure 5H). We calculated correlations between (i) BRD4 protein versus mRNA levels of key partners of fatty acid synthesis pathway (SREBF1, FASN, SCD), (ii) SREBF1 mRNA versus SCD and FASN mRNA, and (iii) SCD and FASN enzymes versus p-RTK (HER2, EGFR, MET) to capture the associations between these molecules. In all cancer types, BRD4 protein levels negatively correlated with SREBF1, SCD and FASN mRNA levels (−0.43 < R < −0.05). This was consistent with the increased expression of fatty acid synthesis genes with BET inhibition we reported above. SREBF1, for which no proteomic data was available, was quantified based on mRNA levels. Consistent with prior reports on the role of SREBF1 on the transcription of lipid metabolism genes (32), SREBF1 and mRNA levels of the two key enzymes, FASN and SCD were strongly correlated (0.28 < R < 0.59). We analyzed RTK activation using available p-RTK measurements (i.e., p-HER2, p-EGFR and p-MET). The relation between SCD and RTK phosphorylation was context-dependent: In HER2+ cells, SCD was highly correlated with p-HER2 (R=0.42) but not with p-EGFR or p-MET. In basal cancers, in which HER2 is not a driver, the correlation between SCD and p-HER2 was reversed (R = −0.22), while a strong correlation between SCD and p-MET was observed (R = 0.47). In ovarian cancers, no p-RTK species demonstrated a strong association (0.06 < R < 0.2). We did not observe similar associations between BRD4 and fatty acid metabolism in luminal A and B breast tumors. Thus, TCGA analysis suggests that the mechanistic link between BRD4, fatty acid metabolism and RTK activation is relevant across a large fraction of HER2-amplified and basal breast cancers.

Finally, we profiled BETi induced phenotypes associated with transcriptomic and proteomic responses in resistant lines. In HCC1954 but not in SKOV3, BETi treatment (48 hours,1μM JQ1) induced morphology changes involving a transformation from a less structured and spread form to a more ordered and contracted state (Figure 6A). Wound scratch assays in monolayer cultures showed that BET inhibition (1μM JQ1) increased cell motility in BETi resistant cells (HCC1954) but not in sensitive cells (SKOV3) (Figure 6B). Consistent with BETi induced SCD upregulation, HCC1954 membrane fluidity increased upon JQ1 treatment (48 hours,1μM JQ1) as quantified by the ratio of monomer to excimer fluorescence of lipophilic pyrene probes, which undergo excimer formation with increasing membrane fluidity. SKOV3 membrane fluidity decreased in response to BETi, consistent with the observed cell motility decrease. As expected, SCDi decreased membrane fluidity in both HCC1954 and SKOV3. We monitored EGFR/HER2 localization changes after JQ1 and SCDi treatment for 48 hours in HCC1954 and SKOV3 cells (Figure 6D). In drug-resistant HCC1954 cells, EGFR/HER2 protein molecules were upregulated and polarized on the cell membrane after JQ1 treatment, leading to an increased local concentration of EGFR/HER2 and likely downstream signaling. No change in HER2 localization was observed in BETi sensitive cells (SKOV3) upon BETi treatment. Thus, BET inhibition induces membrane remodeling including cell morphology, motility, fluidity and receptor tyrosine kinase localization changes associated and consistent with the observed molecular changes in drug resistant cells.

**Figure 6.**
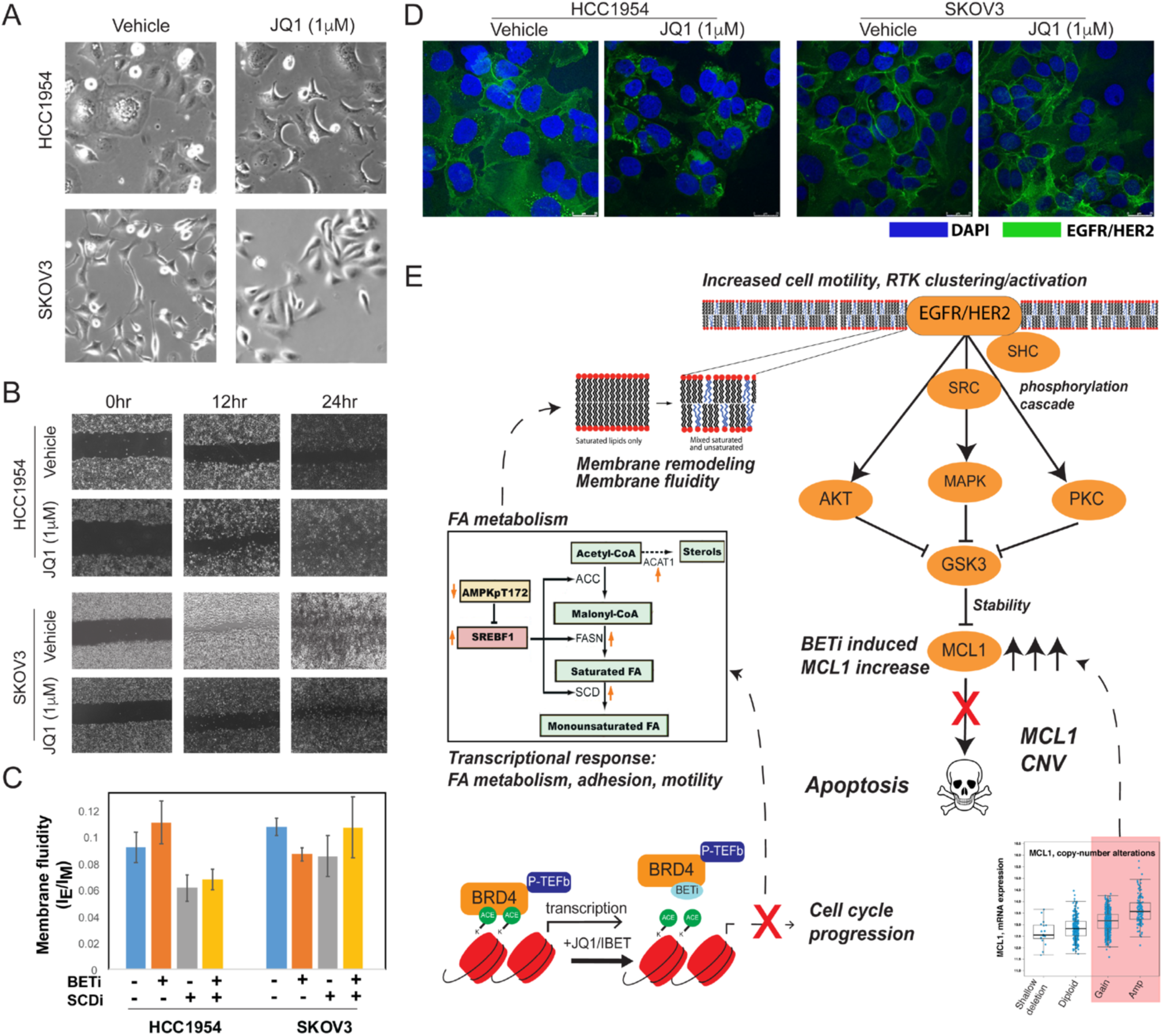
Phenotypic responses to BET inhibition in resistant and sensitive cells. **A.** representative images (20X, brightfield) of cellular morphology changes in response to BETi in JQ1 resistant (HCC1954) and sensitive (SKOV3) cells. **B**. The BETi induces increased motility of HCC1954 (top) and no substantial difference in SKOV3 (bottom) as monitored across time points with a wound scratching assay. **C.** Drug-induced changes in membrane fluidity in HCC1954 and SKOV3 cells based on fluorescent lipophilic pyrene probes that are enriched in excimer state (emission l = 470nm) with increasing membrane fluidity and enriched in monomers (emission l = 400nm) with decreasing membrane fluidity. The membrane fluidity is increased with JQ1 treatment in HCC1954 and decreased in SKOV3. SCD inhibition decreases membrane fluidity as expected. **D.** The representative images of HER2 localization changes in response to BET inhibition in HCC1954 and SKOV3 cells. Images are collected using fluorescent confocal microscopy. BETi induced increased and polarized fluorescence signals from EGFR/HER2 (green) in HCC1954. Whereas the EGFR/HER2 signal was unchanged in response to therapy in SKOV3 cells. **E.** The proposed mechanism of BET induced vulnerability to combined BET and MCL1 targeting in the context of MCL1 chromosomal amplifications or gains based on the integrated multi-omics, network and perturbation analyses. The marked genes and proteins are identified and validated in molecular perturbation response analyses.

## Discussion

We demonstrate that BETis induce a cascade consisting of increases in lipid metabolism mediators, membrane fluidity, cell adhesion and motility. This adaptive response program subsequently leads to the activation of HER2/EGFR signaling and MCL1 upregulation. MCL1 accumulation confers resistance to BET inhibition resulting in synergistic responses to combined inhibition of BET and MCL1 particularly in cells with amplified MCL1. These observations were enabled by a generalizable strategy, integrating high-throughput omics profiling, perturbation biology, network modeling, genomics and functional studies.

Experimental validation in a panel of breast cancer cell lines coupled with genomic analyses in large patient cohorts identified the MCL1 level as a predictor for response to combined inhibition of MCL1 and BETis. MCL1 amplification is a well-characterized prognostic factor in breast cancer (12, 33). Here we showed that MCL1i were effective in cells with MCL1 gain or amplification and that breast cancer cells are further sensitized to MCL1 inhibition when combined with BETis. From a translational perspective, this is particularly important because, MCL1 high-level amplifications are observed in 15% of breast cancer patients, and the frequency of low-level gains in MCL1 is 57%. Therefore, in total, 72% of breast cancer patients fall within the potential candidate cohort for MCL1 targeting as a single agent or in combination with BETi. MCL1 amplification is particularly enriched in the aggressive basal subtype (32%), for which effective therapy options are sorely needed.

Since MCL1 function is also affected by upstream signaling (e.g., GSK3-β) and due to high turnover rates of the protein (34-35), we also studied the signaling mechanisms that may contribute to accumulation of MCL1. Target Score analysis suggested that growth and survival as well as fatty acid pathways were co-upregulated with MCL1 in response to BETi. Guided by the computational analyses and through perturbation-biology-based mechanistic studies and monitoring BETi induced phenotypes, we identified a multi-step mechanism that couples BET inhibition to apoptosis (Figure 6E). In resistant cells, BET inhibition upregulates a cell adhesion, motility and fatty acid synthesis metabolic program, which includes the upstream transcription factor SREBF1 (36) and key enzymes in fatty acid metabolism such as Stearoyl-CoA desaturase (SCD), which converts saturated fatty acids into monounsaturated fatty acids and thus modulates membrane fluidity (37-38). SCD has been suggested as a key modulator of breast cancer cell motility and survival as well as EGFR activation in lung cancer (39-40). Molecular analysis of SCD perturbations suggests that SCD activates HER2/EGFR signaling through membrane remodeling and increased cell membrane fluidity. The resulting spatial changes and polarization of EGFR on the cell membrane are accompanied with increased EGFR phosphorylation and downstream pathway activation consistent with previous reports (41-42). In turn, we show that HER2/EGFR and downstream signaling activation lead to increased MCL1 protein levels. However, no individual signaling pathway activity is obligatory to drive MCL1 accumulation. For MCL1 upregulation, a combined effect from multiple upstream pathways (AKT/PI3K, RAF/MEK, MAPK14, PKC) converging on GSK3 is necessary as demonstrated by pathway inactivation and reversal of adaptive responses with cocktails of up to four drugs. When SCDi and BETi are applied in combination, SCDi reverses the impact of BETi on both HER2/EGFR activation and MCL1 levels. Our integrated analyses suggested concerted roles for MCL1 copy number-driven gene/protein expression and BETi induced signaling plasticity. Together they maintain high levels of MCL1 to evade apoptosis and confer resistance to BET inhibition. In summary, we demonstrated that an interplay between cellular plasticity in response to BET inhibition and copy-number alterations in MCL1 creates a vulnerability to combinations of BET and MCL1 inhibitors. The complementarity and interplay between hard-coded genetic structure and drug-induced plasticity may provide a solution to the long-standing conundrum on the relative dominance of genomics versus signaling activity in driving resistance to targeted agents (Yaffe, 2019). Correlative analyses suggest that the observed mechanisms involving BET, fatty acid synthesis and RTK signaling are also represented in the large TCGA breast cancer patient cohort. Therefore, the observed drug responses are likely to be extrapolatable to clinical settings.

Consistent with previous reports (43-44), Target Score analysis identified a cell cycle arrest signature as a generalized consequence of BET inhibition. In BETi sensitive lines such as SKOV3, apoptosis accompanies cell cycle arrest. Whereas, in resistant cells, a BETi induced transcriptional and signaling program drives evasion of cell death through MCL1 upregulation. Interestingly, in resistant cells, increased cell motility accompanies apoptotic evasion. A compelling speculation is that when under stress, resistant cells not only escape apoptosis but also display a tendency to depart from the cellular locations that mediate stress, potentially reflecting processes inherited through phylogeny.

Other preclinical studies have demonstrated the role of lipid metabolism and cell state plasticity in conferring therapy resistance. For example, high mesenchymal cell states are resistant to diverse therapies, at least in part, stemming from their dependence on lipid peroxidase pathways, and as a result converging on the actionable GPX4 enzyme (45). The compensatory and alternating roles of SCD and FADS2 enzymes in regulating fatty acid desaturation are further evidence for the interplay between cellular plasticity and lipid metabolism (46). Here, for the first time, we linked epigenetic targeting to cellular and signaling plasticity and their consequential drug escape, which can be overcome with the combination of BET and MCL1 inhibition.

Advancing preclinical discoveries to clinical strategies requires a thorough understanding of biomarkers that can identify patients most likely to benefit, mechanisms of drug action and evidence for generalizability. Genomic profiling has already led to major clinical successes in directing subtype-specific treatments such as for HER2-amplified breast cancers (47), BRCA-mutated ovarian and breast cancers (48) or BRAF-mutated melanomas (49). Despite this conceptual revolution and advances in sequencing technologies, durable responses to genetically matched targeted therapies in eligible patient cohorts are limited. As of 2018, only 15% of cancer patients are eligible for a genomics-informed therapy and only 8% of patients benefit from genomics-informed therapy with objective responses (50). The relatively low success rate can be attributed to intra- and inter-tumor heterogeneity, limited genomic profiling techniques, poor genotype-therapy matching criteria, and access to effective drugs. This has led to a strong argument that drug response is not only a linear function of hard-coded genomic aberrations but also mediated by signaling pathway activities, cellular states, and plasticity. This argument implies that clinical decision making needs to be supported with diverse molecular profiling methods including proteomics and RNA expression. Yet, robust molecular proteomic and cellular profiling methods that can diversify decision making in precision medicine and improve patient benefit are still lacking. Combined with clinical studies involving analysis of temporal biopsies from patients under therapy, we argue that this computational/experimental study is a prototype for future clinical applications, which will induce durable responses based on combination therapy decisions that are dynamically informed by multi-omic profiles of tumors evolving under cancer therapy.

## Methods

### Experimental Methods

#### Cell lines and cell culture

HCC1954, HCC1937, HCC1419, HCC70, BT20, BT474, SKOV3, SKBR3, MDAMB468 and MCF7 cell lines were obtained from MD Anderson Cancer Center Cell Line Repository and were thawed two weeks before experiments. There were less than 10 passages between thawing of the cells and the experiments described in this study. In general, mycoplasma testing is performed approximately every 3 months on all cell lines used in the laboratory. MCF-7 cells were maintained in DMEM supplemented with 10% FBS plus antibiotic/antimycotic solution (100U/ml streptomycin and 100U/ml penicillin) (all from Invitrogen). BT20 were maintained in EMEM (Eagle’s Minimum Essential Medium) supplemented with 10% FBS. HCC1954, HCC1937, HCC1419, HCC70, SKOV3, SKBR3 and BT474 were maintained in RPMI 1640 supplemented with 10% FBS. All cells were cultured at 37°C in a humidified atmosphere containing 5% CO_2_.

#### Kinase inhibitors and antibodies

The kinase inhibitors used in this study were purchased from Sellekchem (JQ1 and the inhibitors of EGFR, AKT, PKC, MAPK, P38), ChemiEtek (MCL1 inhibitor-S63845) and MedChemExpress (JQ1 and SCD-inhibitor-A939572). Specific antibodies against MCL1, BCL2, BCL-XL, BRD4, cleaved PARP, p-AKT(T308), p-AKT (S473), p-PKC, p-P38, p-MEK1/2, p-GSK3, and p-HER2 (T1248)/p-EGFR (Tyr1173) were purchased from Cell Signaling (Cell Signaling Technology, MA). Antibody against β-actin and CellBrite Cytoplasmic Membrane Dyes were from Sigma-Aldrich (St. Louis, MO) and Biotium, respectively. Anti-hERBB2/Her2 AF488 antibody was purchased from R&D systems Biotechne.

#### RPPA (Reverse Phase Protein Array)

The cells were washed 3 times with cold PBS and then suspended in RIPA buffer supplemented with proteinase inhibitor and phosphatase inhibitor (Pierce, Rockford, IL, USA). The cell suspension was vortexed for 15 seconds, placed on an end-over-end rotator for 30 min at 4°C and centrifuged at 14,000 x g for 15 min at 4°C. The lysates were prepared to provide 1-1.5mg/ml of total protein lysate. RPPA analysis samples were prepared by adding SDS Sample Buffer, β-mercaptoethanol and RPPA Working Solution to obtain a final concentration of 0.5mg/ml. Samples were heated for 8 min at 100°C, centrifuged 2 min at 14,000 x g and stored at −80°C. The RPPA was performed at the MD Anderson Cancer Center Functional Proteomics core facility.

#### Immunofluorescence imaging experiments

SKOV3 and HCC1954 cells were seeded in coverslips overnight and treated with either vehicle or kinase inhibitors for 48 hours. The cells were fixed with cross-linking method. Cells were switched to fresh growth media with 4% paraformaldehyde for 5 minutes. After 2 times washing with PBS, the cells were incubated in 4% paraformaldehyde (in PBS) for 15 minutes. The fixed cells were rinsed with PBS to remove any fixation agent. Cells were then blocked with odyssey PBS for 1 hour at room temperature and incubated with the anti-hERBB2/Her2 AF488 antibody (1:200, R&D systems biotechne) overnight at 4 °C. After washing with PBS, coverslips were mounted on slides using Prolong gold antifade mountant with DAPI (ThermoFisher Scientific, P36935). Immunofluorescence images were acquired with a confocal microscope (Olympus).

#### Immunoblotting

HCC1954, BT474 and SKOV3 cells were lysed in RIPA buffer (50mM Tris-HCl pH7.4, 150mM NaCl, 1mM EDTA, 1% D.O.C. (Na), 0,1% SDS, 1% Triton X-100) containing protease inhibitors (Pierce, Rockford IL). Protein concentrations were determined using BCA protein assay kit (Pierce, Rockford, IL). An equal amount of proteins was loaded into SDS-PAGE gels and transferred to PVDF membranes (Millipore, MA). Membranes were blocked for 2 hours with 5% non-fat milk in PBST (0.1% Tween-20 in PBS), and incubated overnight with specific primary antibody. After incubation of 2 hours with horseradish peroxidase-conjugated secondary antibody, the proteins were detected by Clarity Western ECL substrate (Bio-Rad).

#### Cell proliferation assays

Cell proliferation was measured by the PrestoBlue Cell Viability Assay kit (A13261, Life Technologies) according to the manufacturer’s instructions. 2∼4×10^3^ cells were seeded into 96-well plates and cultured overnight. After treating with either vehicle or kinase inhibitors for 72 hours, cells were collected and evaluated with a SYNERGY H1 microplate reader with Gen5 software (BioTek). In the viability assays, points and bars represent the mean of triplicates ± SEM. The drug response is quantified as area under the curve by treating the dose-response area as a sum of the trapezoids generated by the responses to consecutive drug doses. The maximum drug dose is quantified as A_max_ = 1-V_max_, where A is the effect at maximum dose and V is the observed viability at maximum dose. The drug synergy is quantified using the Bliss score with the formula; *CI= E*_*AB*_*/(E*_*A*_*-E*_*A*_*(1-E*_*B*_)), where E is the effect to drugs A, B and AB (the combination of drugs A and B).

#### Flow cytometric analysis of cell cycle and apoptosis

2.5×10^5^ of cells (HCC1954, BT474 and SKOV3) were seeded in 6-cm plates. Following treating with vehicle or kinase inhibitors for 48h, cells were harvested, fixed in 1% (W/V) paraformaldehyde in PBS, washed and rehydrated in PBS. DNA was stained according to the instruction of the APO-BRDU™ Kit, by treating the cells with DNA labeling solution, FITC labeled anti-BrdU antibody solution and PI/RNase staining buffer. Flow cytometry was analyzed using BD LSR II flow cytometer and FACSDIV 8.1 software (BD Biosciences) at the MDACC flow cytometry core facility.

#### Migration assays

The effect of drugs on cells’ migratory behavior was analyzed with the scratch wound assay. 4×10^5^ HCC1954 and SKOV3 cells were seeded in 35 mm dishes. After 24 hours, cells were treated with vehicle or kinase inhibitors for 72 hours. The cells were scratched using a sterile 200-μl micropipette tip to form a straight wound. The cells were washed twice with PBS and cultured for an additional 24 hours. The wound closure was measured under an EVOS FL Auto microscope (Life technologies). Images of 3 random fields were acquired at the time point of 0, 6, 12 and 24 h after wounding. The distances traveled by the cells were measured from control and experimental samples and were calculated and compared with time 0.

#### Membrane fluidity

Membrane fluidity was measured using the Membrane Fluidity Kit (Abcam ab189819). After a lipophilic pyrene probe incorporation into the membrane, the monomeric pyrene probe undergoes excimer formation dramatically shifting the emission spectrum of the pyrene probe to a longer red wavelength. The ratio of excimer (emission at 470 nm) to monomer (emission at 400 nm) fluorescence represents a quantitative change of the membrane fluidity. Cells were seeded in 96 well plates, treated with JQ1, SCD inhibitor and JQ1 plus SCD inhibitor for 72 hours. The cells were labeled with labeling solution (15 μM of pyrenedecanoic acid (PDA), 0.08% F-127, in Perfusion Buffer) in the dark for 20 min rocking at 25 °C, washed 2x with Perfusion Buffer, supplemented with culture medium and measured for fluorescence at two wavelengths (excitation at 360 nm, emission at 400 nm and 470 nm) in biological triplicates.

#### mRNA Sequencing

Cells (HCC1954, SKOV3 and BT474) were collected after treating with 1 μM of BETi (JQ1) for 24 hours all in biological duplicates. Total RNA was isolated and purified using the RNeasy Plus Mini Kit (Qiagen). After quantification with Qubit 2.0 (Life Technologies), the samples passed through quality control steps for sample integrity and purity with Agilent 2100 and Agarose Gel Electrophoresis. After the sample QC, the cDNA library is constructed using the NEBNext Ultra II RNA Library Prep Kit (NEB) as follows: mRNA is enriched using oligo(dT) beads. The mRNA is then fragmented via sonication in fragmentation buffer, followed by cDNA synthesis using random hexamers and reverse transcriptase. After first-strand synthesis, a custom second strand synthesis buffer and enzyme mix are added to generate the second strand by nick translation. The final cDNA library is ready after a round of DNA purification, end-repair, A-tailing, ligation of sequencing adapters, size selection and PCR enrichment. For library QC, library concentration was quantified using a Qubit 2.0 fluorometer (Life Technologies) and then diluted to 1 ng/μl before checking insert size on an Agilent 2100. Libraries were then quantified to greater accuracy by quantitative PCR (Q-PCR) prior to sequencing. RNA libraries were sequenced with 20M paired-end 150bp reads on the Illumina NovaSeq 6000 platform (Novogene Inc.). Raw data was subjected to a round of QC to remove low-quality reads. The resulting mRNA sequence count data was normalized to enable cross-sample analyses.

#### Breast cancer xenograft models

Nude athymic NCr female mice (5 weeks old) were obtained from the Department of Experimental Radiation Oncology, MD Anderson Cancer Center. All studies were conducted according to the experimental protocol approved by the MD Anderson Institutional Animal Care and Use Committee. MDAMB468 cells (2×10^6^ in 20% matrigel) were orthotopically injected into the mammary fat pad of each mouse. After approximately 2 weeks, when tumor sizes reach 3-5 mm, mice were randomized in the treatment groups, including empty liposomes, liposomal JQ1, MCL1 inhibitor (S83445) or combination groups. Treatments were administered at 20 mg/kg dose by i.p. injection into mice (9 mice/per group, 10 in control arm) 3 times/week. Liposomes were prepared based on our previously published method. Two animals from each arm were sacrificed at day 10 and the terminal time point for IHC analyisis of response markers on therapy.

#### Preparation of liposomal nanoparticles

Liposomal nanocarriers were prepared with Dimyristoyl-sn-glycero-3-phosphocholine (DMPC) and pegylated distreroly-phosphotidyl ethanolamine (DSPE-PEG-2000) (Avanti Lipids) as previously described by us (51). Briefly, DMPC and DSPE-PEG2000 were mixed at the ratio of (10:1) and were mixed with small molecule inhibitors at a ratio of 10:1 (w/w) and lyophilized in the presence of excess tertiary butanol. After liposomal drugs were reconstituted in PBS, the agents were systemically administered to mice.

#### Immunohistochemistry

At the termination of the xenograft experiment, animals were euthanized according to guidelines by the Institutional Animal Care and Use Committee, adopted from the AVMA Guidelines for the Euthanasia of Animals: 2013 Edition. The mice tissue samples were fixed in 10% neutral buffered formalin for 24 hours and transferred to 70% ethanol until ready to be processed. Samples were embedded in paraffin, sectioned at 5 mm and stained routinely with hematoxylin and eosin. Unstained sections were designated for immunohistochemical staining. BCL2 (Cell Signaling. Catalog No. 15071), SCD (Cell Signaling. Catalog No. #2794), Clvd-Caspase3 (Cell Signaling. Catalog No. 9661), KI67 (Abcam. Catalog No. ab15580), BRD4 (EMD Millipore. Catalog No. ABE1391) and MCL1 (Cell Signaling. Catalog No. 39224) immunoreactivity was detected using manual staining with the R.T.U. Vectastain Universal Elite ABC kit Anti-mouse IgG/Rabbit IgG (Vector Laboratories. Catalog No. K-7100). Heat-induced antigen retrieval was done using a pH 6 Citrate-based buffer (made in house) and a tris-EDTA pH 9.0 buffer (made in house) for 30 minutes in the IHC-TekTM Epitope Retrieval Steamer (IHC World. Catalog No. IW-1102). The primary mouse monoclonal anti-BCL2 antibody was incubated at 1:400 dilution, the primary rabbit monoclonal anti-SCD was incubated at 1:200 dilution, the primary rabbit polyclonal anti-Clvd-Caspase3 was incubated at 1:200 dilution, the primary rabbit polyclonal anti-KI67 was incubated at 1:300 dilution, the primary rabbit polyclonal anti-BRD4 was incubated at 1:250 dilution, and the primary rabbit monoclonal anti-MCL1 at 1:100 dilution for one hour at room temperature in an incubation chamber. Immunoreactivity was detected using DAB (diaminobenzidine) Peroxidase (HRP) Substrate Kit (Vector Laboratories, Catalog No. SK-4100) incubated for approximately 2 minutes. The anti-rabbit/mouse biotinylated universal antibody was applied for 30 minutes at room temperature. The Vectastain ABC reagent was incubated for 30 minutes at room temperature. Slides were counterstained with Mayer’s hematoxylin (Electron Microscopy Sciences. Cat No. 26173-03) for 3 minutes. Hematoxylin was enhanced with 0.25% ammonium hydroxide bluing solution (made in house) for 1 minute at room temperature. Sections of the experimental control group served as positive controls for validation. Negative control slides were stained by substituting the primary antibody with 2.5% normal horse serum.

### Computational Methods

#### Target Score algorithm

The aim of the network-level description of adaptive responses is to better explain collective mechanisms, in contrast with conventional analysis which focuses on individual genes/proteins. The algorithm: (i) quantifies and visualizes collective adaptive responses to targeted therapy; (ii) nominates combination therapies, involving the agent that induces the adaptive response and a second agent that targets vulnerabilities induced by the adaptive response (see methods). The algorithm is formulated such that highest target scores correspond to potential adaptive responses (drug-induced activation of oncogenic processes or deactivation of tumor suppressors). Phosphoproteins with the lowest target scores correspond to direct response mechanisms to targeted perturbations. The Target Score algorithm nominates modules of functionally related molecular entities involved in adaptive resistance with a higher likelihood of therapeutic relevance. Target Score also reduces the number of potential false positives as it eliminates signals from proteins with no connection to collective processes. The Target Score algorithm works in multi-steps involving the construction of a reference network, target score calculations, statistical assessment of scores, and the determination and visualization of response pathways.

### Code availability

Target Score code is available as the Docker container cannin/targetscore:mcl1-analysis with additional instructions here: https://hub.docker.com/repository/docker/cannin/targetscore

### Input data and data quality

The molecular response data collected before and after therapy is used as the input in the analysis pipeline. The BETi RPPA response data is generated at the MDACC functional proteomics core facility. The functional proteomics core validates each antibody by analysis of western blots to observe each readout as a single or dominant band on a blot and ensure the Pearson correlation coefficient between RPPA and western blot readouts are greater than 0.7. The dynamic range and specificity are routinely determined using peptides, phosphopeptides, growth factors, Inhibitors, RNAi, cells with wide levels of expression including 330 cell lines under multiple conditions on a single array. The intra and inter-slide reproducibility of RPPA reads are routinely monitored. In this study, BETi responses from cell lines were interrogated using 217 antibodies at time points of 24 and 48 hours after drug treatment in cells cultured as 2D monolayers and 3D matrigel cultures. However, the method can analyze data from both patient biopsy samples and cell lines. The method is amenable to analysis of drug response data from a single sample to detect drug-induced vulnerabilities in a sample-specific way or thousands of conditions to determine complex statistical patterns of adaptive responses across varying genomic contexts or perturbation agents. The analysis can be performed with time series (short term perturbation periods or long term acquired resistance events) or multi-dose perturbation data to monitor the evolution of adaptive response patterns in time and dose space.

### Reference network construction

Network models are constructed using the signaling data stored in Pathway Commons (PC) database and manual curation/correction by experts. PC integrates detailed human pathway data from multiple public resources such as Reactome, NCI PID, PhosphoSitePlus, Panther Pathways, and HumanCyc (52). Automated prior pathway extraction is achieved using the SignedPC module embedded in the BioPAX/PaxTools pathway analysis language and CausalPath method (53-55). In order to delineate the pathway interactions captured by the RPPA data, we identified 4 different relation types: phosphorylation, dephosphorylation, expression upregulation, and expression downregulation. The first two relations can be used to explain phospho-protein changes, and the last two can be used to explain total protein or gene expression changes. Using the BioPAX-pattern framework (56), we define a set of patterns to detect such relations and identify the signaling interactions between the phosphoproteomic entities. SignedPC resource includes 130158 interactions, of which 107251 define “expression upregulation”, 20964 define “phosphorylation”, 1942 define “expression downregulation” and 2660 define dephosphorylation events. The details of the construction of SignedPC is described in (55). The resulting interaction set extracted from the SignedPC forms the reference network.

### The Target score and target detection

A target score that quantifies the adaptive pathway responses to a perturbation as a sum of the response from each individual protein and its pathway neighborhood is calculated for each entity on the reference network. The calculation combines the cell type-specific drug response data with signaling network information extracted from signaling databases. High target score identifies proteins involved in adaptive response (e.g., upregulation of receptor tyrosine kinase expression by MEK inhibitor via a feedback loop^4^) and low target score corresponds to the immediate impact of the drug (e.g., inhibition of ERK phosphorylation by MEK inhibitor). The mathematical formulation of the Target Score is

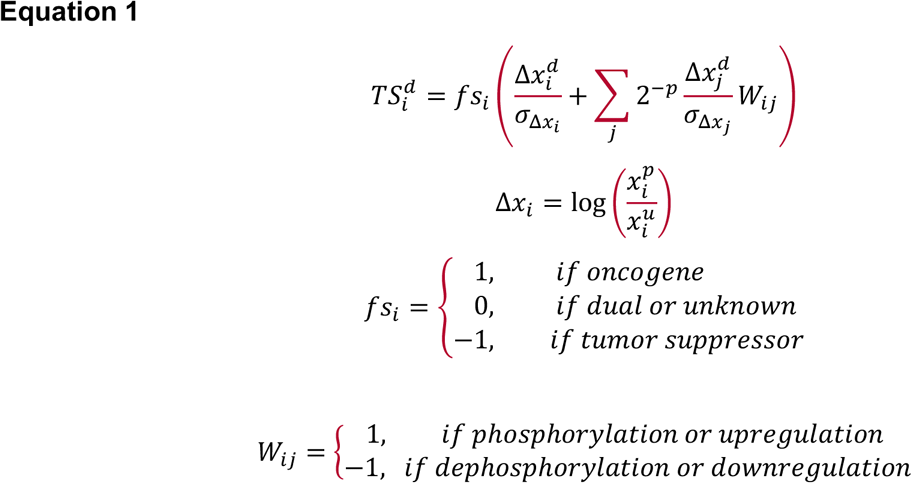

where *fs*_*i*_ represents the functional score (see below) of the node of interest, *Δx*_*i*_ is the proteomic response, log normalized with respect to the unperturbed (or pretreatment) level. The *σ* is the standard deviation of *Δx*_*i*_ over all perturbation conditions and all drug doses for a sample. *fs*_*i*_ scores ensure high positive target scores correspond to adaptive responses. The division of *Δx* by *σ* normalizes a readout with respect to the dynamic range of the corresponding antibody. Node j is a node in the pathway neighborhood of the node i, with readout *Δx*_*j*_ and standard deviation. *p*_*ij*_ is the pathway distance between nodes i and j. W_ij_ represents the signaling interaction between nodes i and j extracted algorithmically from databases (i.e., reference network). The cumulative target score over multiple doses can be calculated as the area under the curve of the target score at each dose versus drug doses.

#### A functional score (*fs*)

is assigned to each of the proteomic entities measured in the MDACC RPPA and DFCI/MSKCC Zeptosens proteomics platforms (57-58). We used manual curation, resources such as Phosphosite database and TUSON tumor suppressor/oncogene resource (59) results. We assigned (+1) for total levels or activating phosphorylation of oncoproteins and inhibitory phosphorylation of tumor suppressors. Similarly, a functional score of (−1) is assigned to total levels or activating phosphorylation of tumor suppressors and inhibitory phosphorylation of oncoproteins.

#### The statistical assessment

detects and eliminates the target scores that are likely driven solely by the network structure with no significant impact from the cell type-specific data. For this purpose, the probability of observing a given target score is calculated over a null distribution of target score values generated with randomized drug response data and the fixed network structure. A random sampling of proteomic responses (randomized antibody label mixing) at each drug dose generates randomized data sets over all antibody readouts. The target score is calculated for 1000 independent random datasets using equation 1, and the null score distribution is constructed. Next, the FDR-adjusted P value is calculated for the target score value from actual data. For increased statistical power, the p-values across biologically similar or highly overlapping conditions (e.g., similar BETis) that lead to similar target scores are merged using the Stouffer’s method with the formula (60).

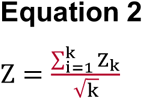

Z scores are obtained through p-value-to-Z conversion and k is the total number of conditions merged. Resulting Z-value is back converted to P-values and FDR-adjusted using the Benjamini-Hochberg method.

#### Determination of response pathways

The resistance and response (drug activity) pathways are detected by the reanalysis of the target scores and the underlying network models. Proteins that are connected within the first neighborhood of each other in the underlying network and have the highest scores (20 top entities) represent the potential adaptive response pathway(s) and those with low target scores represent the potential response/drug activity pathway. The pathway relations are extracted from the reference network model and visualized using the Cytoscape software (61).

### Differential analysis of omics response

The proteomic response was quantified by normalizing the median normalized proteomic readouts from drug-treated samples with respect to the matched untreated samples. The proteomic differential response (Figure 5) is quantified as the S_t,i,_ = ∑_d_[(1/N)*[(*Δ*X^t,d^_HCC1954,i_-*Δ*X^t,d^_SKOV3,i_)+(*Δ*X^t,d^_BT474,i_-*Δ*X^t,d^_SKOV3,i_)], where *Δ*x is the phosphoproteomic response for entity i, t is the time point (24 or 48h), and d represents BETis (JQ1, IBET151, IBET726, or IBET762), N is the number of drugs used (N=4). The statistical significance of proteomic differences is quantified using a paired 2-tailed t-test followed by a Bonferroni multiple hypothesis correction with the null hypothesis Ho: **μ**(X^t,d^_HCC1954,i_)= **μ**(X^t,d^_SKOV3,i_) for readouts across both time points (t) and all BET inhibitors (d) and FDR adjusted across each proteomic measurement (i). Statistical significance is assigned to cases, Q<0.05. The transcriptomic response was quantified by normalizing the mRNA counts from drug-treated samples with respect to the matched untreated samples. The significant mRNA fold changes upon drug treatment were detected as FDR adjusted Q-values < with the DESeq method. To eliminate noise from rare transcripts, we included the mRNA species with an abundance over a minimal threshold (minimum count > 200) in at least one condition (i.e., before or after treatment in at least one cell line). The mRNA fold changes that significantly increased in response to BET inhibition in the resistant lines (HCC1954, SKOV3) and significantly decreased in response to BET inhibition in the sensitive line (SKOV3) were included in the differential analysis. The differential responses are visualized as a heatmap that demonstrates fold differences of mRNA species that change in opposite directions in resistant vs. sensitive lines.

### Go-Term enrichment

We downloaded GO-term gene associations from http://geneontology.org/gene-associations/goa_human.gaf.gz, and tested the enrichment of the terms in differentially expressed 127 genes using Fisher’s exact test. We used the Benjamini-Hochberg method to select the significance threshold to get a result set with 0.1 FDR.

### Analysis of patient and cell line genomics and proteomics data

The correlative analyses of breast and ovarian cancer patients are performed using the TCGA breast invasive carcinoma (N= 1108) and ovarian serous cystadenocarcinoma (N=606) provisional data stored in the cBioPortal. The patients in the basal breast cancer subtype are selected based on the annotation in (62). The HER2 amplified breast cancer cases are selected based on the ERBB2 gene GISTIC copy number assignments stored in the cBioPortal. The correlations between molecular entities are computed using the TCGA microarray data for mRNA and RPPA data for phosphoproteomics. The Pearson’s correlations are computed using the cBioPortal analysis and visualization suits. The cell line genomics analyses are performed using the data from the CCLE resource (26,63). The cell line mutation data is based on WGS and WES followed by a mutation calling for SNVs and InDels, and filtering out germline variants performed at the Broad Institute as described in (63). The copy number data is based on SNP arrays processed with circular binary segmentation of copy number levels and GISTIC copy number calls as described in (26).

## Acknowledgement

This work is supported with grants from MD Anderson Cancer Center Support Grant P30 CA016672 (the Bioinformatics Shared Resource), OCRF Collaborative Research Award, U.S. National Cancer Institute grants U24CA210950, P50CA217685, U01CA217842 Fund for Innovation in Cancer Informatics (ICI), UT STARS Rising Stars Award, CPRIT High-Impact/High-Risk Award (RP170640), and a kind gift from the Miriam and Sheldon Adelson Medical Research Foundation, SAC110052 from the Susan G. Komen Foundation, and Breast Cancer Research Foundation BCRF-18-110. Confocal microscopy experiments are performed at the MDACC Advanced Microscopy Facility funded by NIH S10 RR029552.

