## Supplemental for "Adaptive response to BET inhibition induces therapeutic vulnerability to MCL1 inhibitors in breast cancer"

#### Supplemental Figures

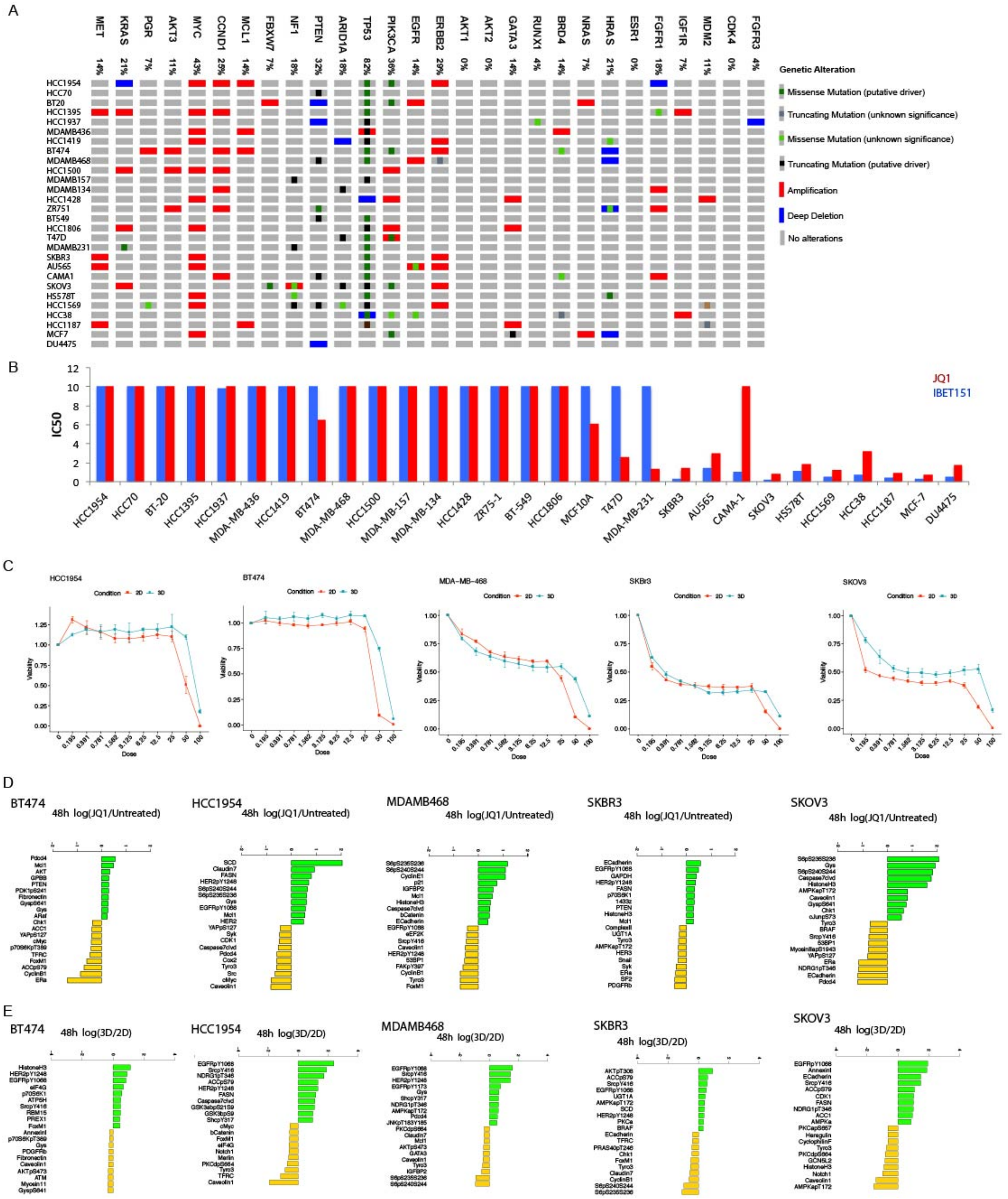


***Figure S1. A.*** *Genomic alteration profiles in the breast cancer and ovarian (SKOV3) cell lines treated with BETis. The order of cell lines reflects BETi sensitivity rank (top: most resistant, bottom: most sensitive). Genes that are frequently altered in breast cancer are included in the analysis.* ***B.*** *BETi (JQ1 and IBET151) cell viability response IC50 for cell lines.* ***C.*** *Dose response curves of BET inhibition in HCC1954, BT474, MDAMB468, SKBR3, and SKOV3 cultured in 2D*

*and 3D.* ***D.*** *The responses to BET inhibition (JQ1 at 48hrs). Substantial changes (X>1 in log2 space) are observed in SCD in HCC1954, phospho-S6 in MDAMB468 and SKOV3, CyclinE1 in MDAMB468, and Gys, HistoneH3 and capspase7 clvd in SKOV3. Note that MCL1 protein is top 10 most upregulated proteins in BT474, HCC1954, MDAMB468 and SKBR3 but it is never the most upregulated protein*. ***E.*** *Comparison of proteomic responses to BETi (JQ1) in 3D cultures vs. 2D cultures in all cell lines. The bar plots display relative values of response to JQ1 with respect to the responses in monolayered cultures. Most substantial increases are in EGFR phosphorylation at Y1068 (up to 2.4 increase in all cell lines except SKOV3), HER2 phosphorylation at Y1248 (log-fold increase of 0.4 in SKBR3 to 1.4 in SKOV3), HistoneH3 levels (1.2 log2-fold increase in BT474) and SRC phosphorylation at Y416 (up to 2 log-fold increase in all cell lines). Most substantial reduction induced by ECM attachment were in S6 phosphorylation at S235/236 and S240/244 (~1 log2-fold decrease in SKBR3 and MDAMB468), caveolin (2 log2-fold decrease in HCC1954) and AMPK_pT172 (1.6 log-fold decrease in SKOV3). AKT phosphorylation at T308 was increased in SKBR3; E-cadherin and annexin were increased in SKOV3 upon ECM attachment. Interestingly, responses in lipid metabolism proteins such as FASN and ACC1 (total and phosphorylated state) were higher in 3D cultured HCC1954 and SKOV3 lines.*

A

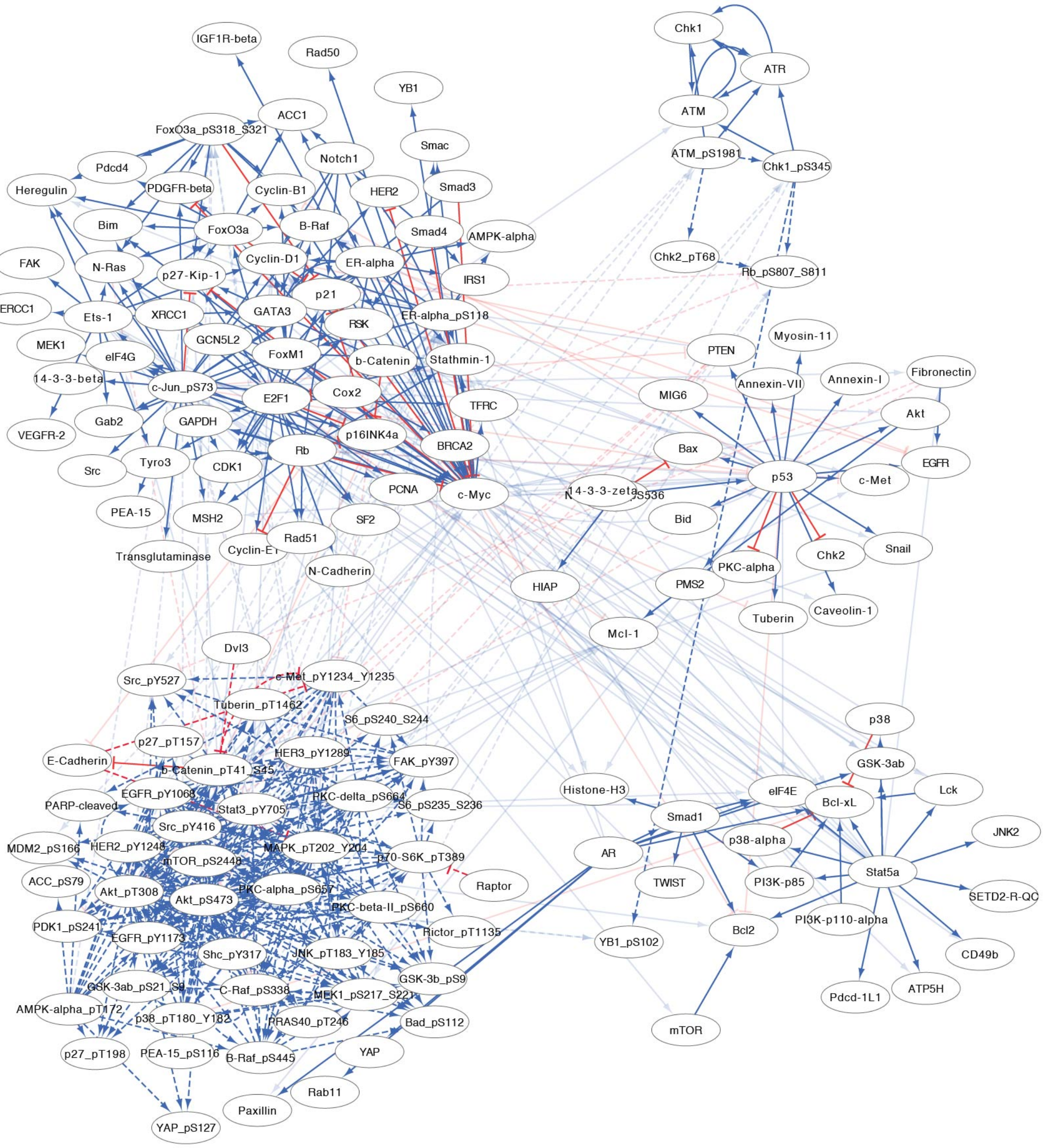

IGF1R-beta
Rad50
YB1
ACC1
FoxO3a_pS318_S321
Smac
Notch1
Pdcd4
Smad3
Heregulin
PDGFR-beta
Cyclin-B1
HER2
Bim
FoxO3a
B-Raf
Smad4
AMPK-alpha
FAK
N-Ras
Cyclin-D1
ER-alpha
p27-Kip-1
IRS1
ERCC1
p21
XRCC1
GATA3
ER-alpha_pS118
Ets-1
MEK1
GCN5L2
RSK
eIF4G
FoxM1
b-Catenin
Stathmin-1
14-3-3-beta
c-Jun_pS73
Cox2
E2F1
TFRC
Gab2
VEGFR-2
GAPDH
p16INK4a
BRCA2
Tyro3
Rb
Src
CDK1
PCNA
c-Myc
PEA-15
MSH2
SF2
Rad51
Transglutaminase
Cyclin-E1
N-Cadherin
Chk1
ATR
ATM
ATM_pS1981
Chk1_pS345
Chk2_pT68
Rb_pS807_S811
Myosin-11
PTEN
Annexin-VII
Annexin-I
Fibronectin
MIG6
Akt
Bax
EGFR
p53
c-Met
14-3-3-zeta
Bid
Snail
PKC-alpha
Chk2
HIAP
PMS2
Tuberin
Caveolin-1
Mcl-1
Dvl3
Src_pY527
c-Met_pY1234_Y1235
Tuberin_pT1462
S6_pS240_S244
p38
p27_pT157
E-Cadherin
HER3_pY1289
FAK_pY397
GSK-3ab
b-Catenin_pT41_S45
Histone-H3
eIF4E
Lck
EGFR_pY1068
PKC-delta_pS664
PARP-cleaved
Stat3_pY705
S6_pS235_S236
Smad1
Bcl-xL
JNK2
Src_pY416
MDM2_pS166
HER2_pY1248
MAPK_pT202_Y204
p38-alpha
AR
mTOR_pS2448
p70-S6K_pT389
Stat5a
ACC_pS79
Raptor
PI3K-p85
TWIST
Akt_pT308
Akt_pS473
PKC-alpha_pS657
SETD2-R-QC
PKC-beta-II_pS660
PDK1_pS241
Rictor_pT1135
PI3K-p110-alpha
Bcl2
YB1_pS102
EGFR_pY1173
Shc_pY317
JNK_pT183_Y185
CD49b
GSK-3b_pS9
GSK-3ab_pS21_S9
AMPK-alpha_pT172
C-Raf_pS338
MEK1_pS217_S221
Pdcd-1L1
ATP5H
p38_pT180_Y182
PRAS40_pT246
Bad_pS112
mTOR
p27_pT198
PEA-15_pS116
B-Raf_pS445
YAP
Rab11
Paxillin
YAP_pS127

B

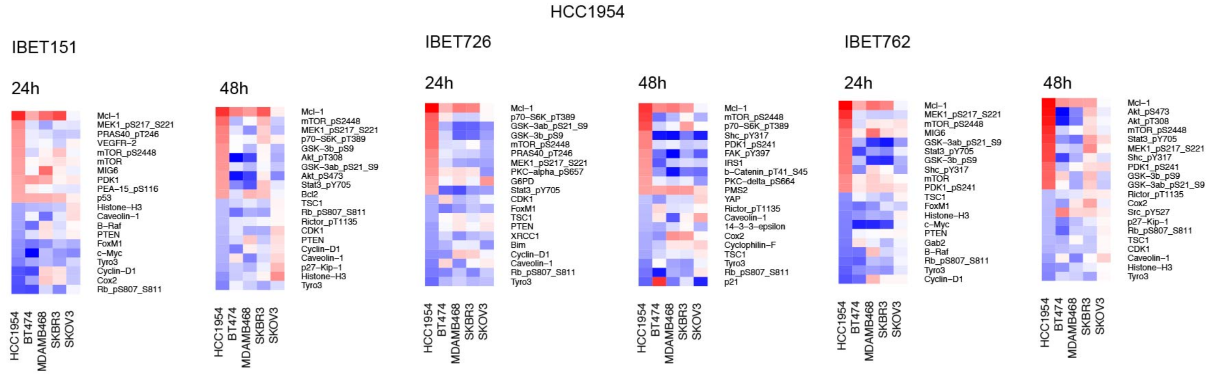
HCC1954
IBET151
IBET726
IBET762
24h
48h
HCC1954
BT474
MDAMB468
SKBR3
SKOV3

C

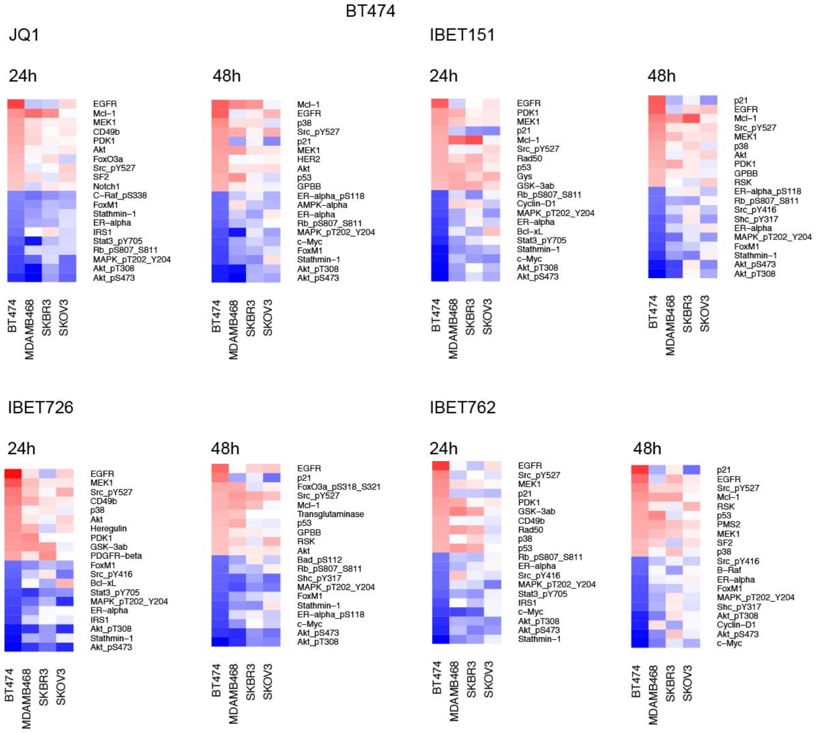
BT474
JQ1
IBET151
IBET726
IBET762
24h
48h
BT474
MDAMB468
SKBR3
SKOV3

D

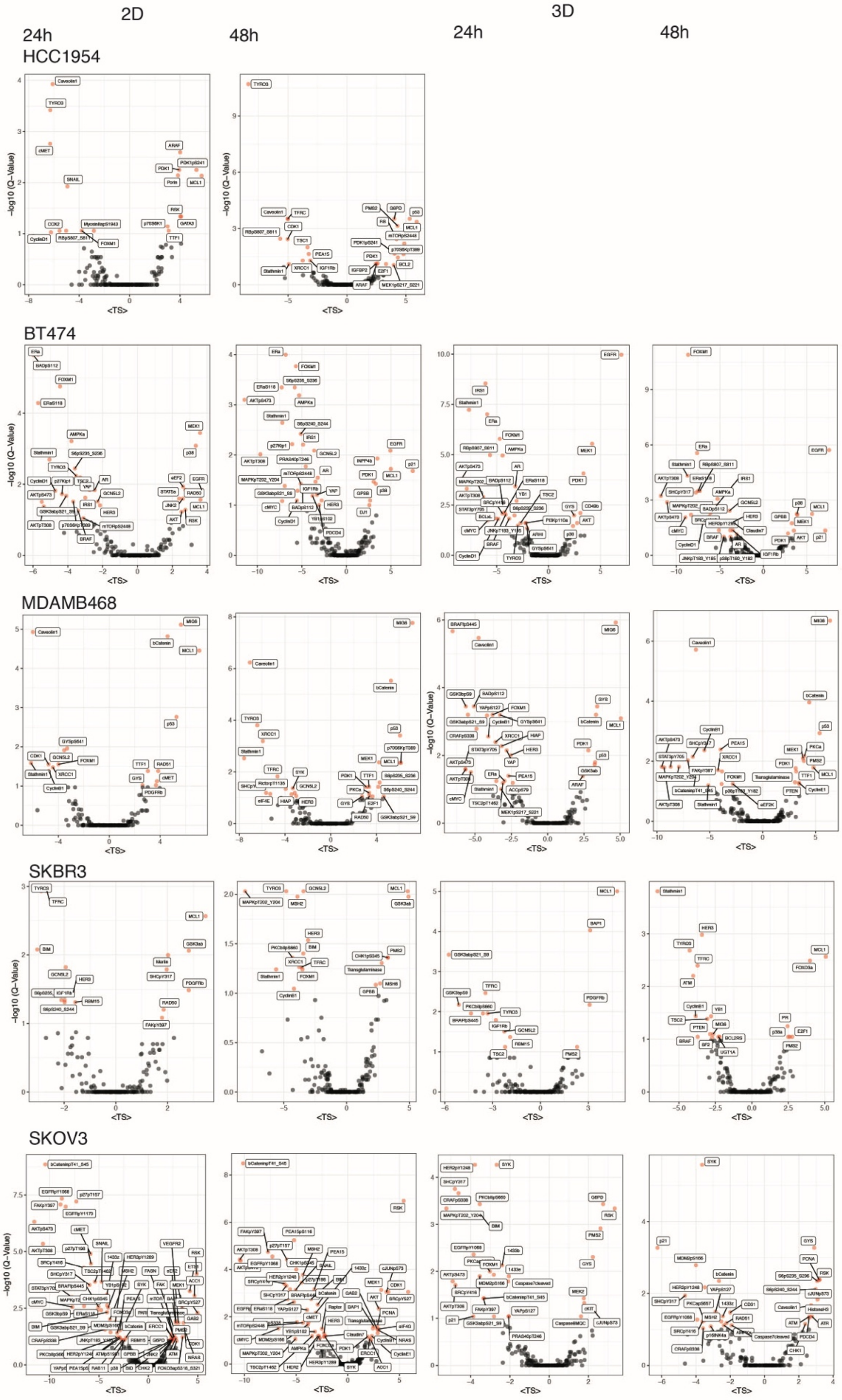
2D
3D
24h
48h
24h
48h
HCC1954
BT474
MDAMB468
SKBR3
SKOV3
-log10 (Q-Value)
<TS>

***Figure S2.*** *Target Score analysis of HCC1954, SKOV3, BT474, MDAMB468 and SKBR3 lines treated with BETis in 2D and 3D cultures based on RPPA data collected 24 and 48 hours post perturbation.* ***A.*** *The reference network used in Target Score calculations.* ***B****. The heatmaps of target scores calculated for each drug treattment in HCC1954 cells cultured in 3D cultures (JQ1 plot is in Figure 2). He cell lines are ordered from most resistant (HCC1954) to most sensitive (SKOV3). The proteins with top10 highest score and bottom10 lowest target scores from HCC1954 are listed. The readouts in more sensitive cell lines are provided for comparison.* ***C.*** *Similar heatmaps as in (B) with based on the top and bottom target scores for BT474 cells.* ***D.*** *The volcano plots list the most significant target scores (Q value on y-axis) and highest average target scores across 4 BET inhibitors (<TS> in x-axis). The results are given for both 2D and 3D cultured cells for all cell lines analyzed at 24h and 48h.*

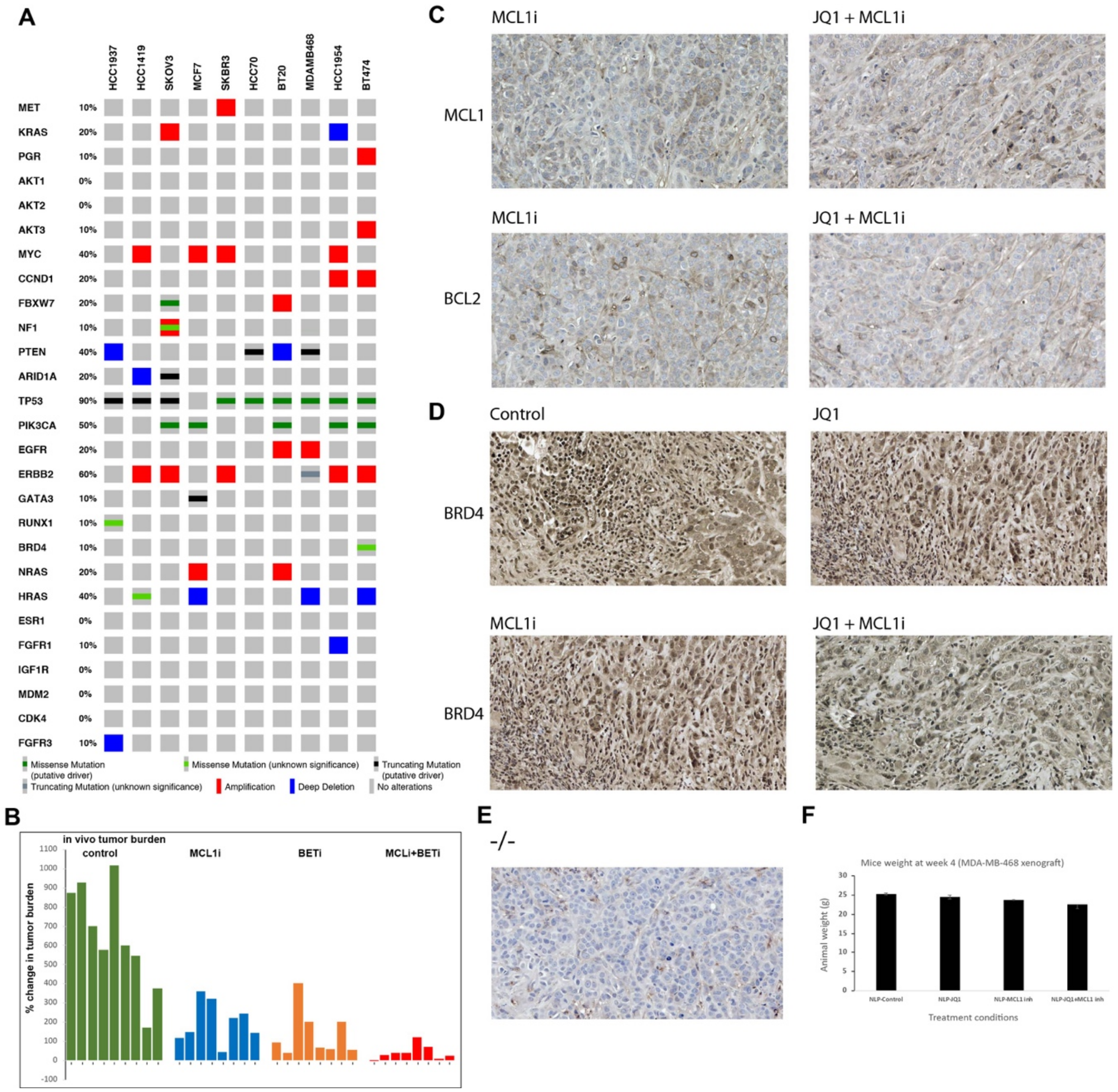


***Figure S4. A.*** *The genomic alterations in the breast and ovarian (SKOV3) cell lines tested with the BET and MCL1 inhibitor combination.* ***B.*** *The tumor burden change in control and drug treatment (BETi, MCL1i, combination) in MDMB468 xenograft model.* ***C.*** *Representative images of immunohistochemical analysis of BRD4 expression in drug treated tumor xenograft models.* ***D.*** *Representative images of immunohistochemical analysis of MCL1 and BCL2 expression in MCL1 inhibitor and of MCL1 and BET inhibitor combination treated tumor xenograft models. N=9/arm for control, N=8/arm for treatment cohort. One animal was sacrificed for IHC analyses at end point and excluded from analysis.* ***E.*** *The negative control IHC staining with no primary antibody treatment.* ***F.*** *Animal weights in each condition at the termination time of the experiment.*

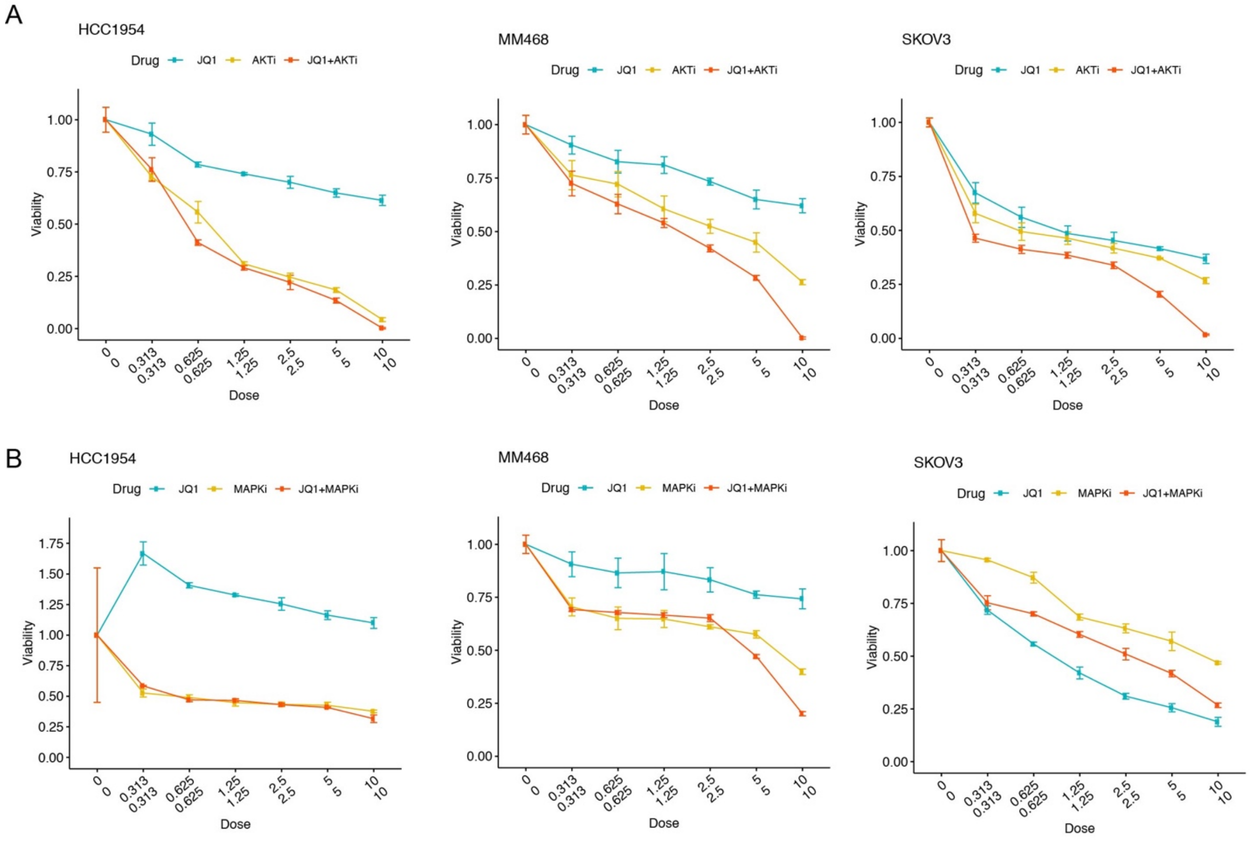


***Figure S5.*** *The cell viability response to combination of BET inhibitors with* ***A****. AKT inhibitor,* ***B.*** *MEK1/2 inhibitors (trametinib).*

### Supplementary tables

***Table 1***. *GO-term enrichment analysis of differential mRNA expression in BETi resistant vs. sensitive cells*

| P-value | GO Term | Hit genes in the query |
|---|---|---|
| **5.51E-06** | GO:0046390 ribose phosphate biosynthetic process | [PFKFB2, AK3, ACAT1, PAICS, RFK, CAD, ELOVL7, SCD, PFKP] |
| **2.52E-05** | GO:0044281 small molecule metabolic process | [CYP1B1, SLC16A3, GPT2, SLC6A14, SLC27A3, PFKFB2, AK3, ACAT1, INPP5J, ASL, ST3GAL1, B4GALNT2, TMEM86A, PAICS, ACSF2, GALK2, RFK, SREBF1, CAD, ELOVL7, CD44, SC5D, PTGES2, SCD, SLC25A32, PFKP] |
| **2.58E-05** | GO:0016020 membrane | [TES, F11R, RIMS4, CDH1, CYP1B1, SLC39A8, PIP4K2C, GYG1, SLC16A3, SLC6A14, KRT8, ALG3, SLC5A6, MAP1B, TRIM13, EPHA1, SLC27A3, LLGL2, SLC41A2, CYB5R1, KLC2, ZMAT3, TSPAN9, INPP5J, ST3GAL1, B4GALNT2, SPECC1, TMEM120B, TMEM86A, ST14, GLMP, L1CAM, PAICS, CD24, MCU, HLA-DRB1, NCAPG2, GRIK3, FMN1, TMEM47, MRPL4, SPRED2, TUBA1A, HRAS, SREBF1, CAD, ELOVL7, TMC6, VPS37B, RHOB, CEACAM6, CLDN12, TMEM106B, CDH13, PGAM5, CD44, SLC29A2, SGMS2, SLC7A1, TRAK2, ERBB3, PALMD, SC5D, MAL2, LRRC8B, PTGES2, GALNT1, SCD, VPS41, F2RL1, PTPN6, SLC25A32, PFKP] |
| **4.09E-05** | GO:0019752 carboxylic acid metabolic process | [CYP1B1, SLC16A3, GPT2, SLC6A14, SLC27A3, PFKFB2, ACAT1, ASL, ST3GAL1, ACSF2, CAD, ELOVL7, CD44, PTGES2, SCD, SLC25A32, PFKP] |
| **6.73E-05** | GO:0009165 nucleotide biosynthetic process | [PFKFB2, AK3, ACAT1, PAICS, RFK, CAD, ELOVL7, SCD, PFKP] |
| **7.51E-05** | GO:1901293 nucleoside phosphate biosynthetic process | [PFKFB2, AK3, ACAT1, PAICS, RFK, CAD, ELOVL7, SCD, PFKP] |
| **1.02E-04** | GO:0009259 ribonucleotide metabolic process | [PFKFB2, AK3, ACAT1, PAICS, ACSF2, RFK, CAD, ELOVL7, SCD, PFKP] |
| **1.32E-04** | GO:0043436 oxoacid metabolic process | [CYP1B1, SLC16A3, GPT2, SLC6A14, SLC27A3, PFKFB2, ACAT1, ASL, ST3GAL1, ACSF2, CAD, ELOVL7, CD44, PTGES2, SCD, SLC25A32, PFKP] |
| **1.54E-04** | GO:1901137 carbohydrate derivative biosynthetic process | [ALG3, PFKFB2, AK3, ACAT1, ST3GAL1, PAICS, RFK, CAD, ELOVL7, SCD, PFKP] |
| **1.55E-04** | GO:0019693 ribose phosphate metabolic process | [PFKFB2, AK3, ACAT1, PAICS, ACSF2, RFK, CAD, ELOVL7, SCD, PFKP] |
| **1.70E-04** | GO:0006082 organic acid metabolic process | [CYP1B1, SLC16A3, GPT2, SLC6A14, SLC27A3, PFKFB2, ACAT1, ASL, ST3GAL1, ACSF2, CAD, ELOVL7, CD44, PTGES2, SCD, SLC25A32, PFKP] |
| **2.31E-04** | GO:0030155 regulation of cell adhesion | [CDH1, CYP1B1, EPHA1, B4GALNT2, BMP7, CD24, FMN1, CEACAM6, CDH13, CD44, SEMA3E, ERBB3, PTPN6] |
| **2.59E-04** | GO:0031224 intrinsic component of membrane | [F11R, EVA1B, CDH1, SLC39A8, SLC16A3, SLC6A14, ALG3, SLC5A6, TRIM13, SIDT1, EPHA1, SLC27A3, SLC41A2, CYB5R1, TSPAN9, ST3GAL1, B4GALNT2, TMEM120B, TMEM86A, ST14, GLMP, L1CAM, CD24, MCU, HLA-DRB1, GRIK3, TMEM181, TMEM47, SREBF1, ELOVL7, TMC6, CEACAM6, CLDN12, TMEM106B, CDH13, PGAM5, CD44, SLC29A2, SEMA3E, SGMS2, SLC7A1, MIEN1, ERBB3, SC5D, MAL2, LRRC8B, TM4SF18, PTGES2, GALNT1, SCD, F2RL1, SLC25A32] |

| | | |
|---|---|---|
| **2.96E-04** | GO:0090407 organophosphate biosynthetic process | [PIP4K2C, PFKFB2, AK3, ACAT1, INPP5J, PAICS, RFK, CAD, ELOVL7, SGMS2, SCD, PFKP] |
| **3.08E-04** | GO:0072522 purine-containing compound biosynthetic process | [PFKFB2, AK3, ACAT1, PAICS, ELOVL7, SCD, PFKP] |
| **3.19E-04** | GO:0006810 transport | [F11R, RIMS4, SLC39A8, GYG1, SLC16A3, PPFIA3, SLC6A14, SLC5A6, MAP1B, SIDT1, SLC27A3, LLGL2, SLC41A2, CYB5R1, KLC2, CRACR2B, ZMAT3, CD24, MCU, GRIK3, TUBA1A, MB, HRAS, TIGD5, CAD, TMC6, VPS37B, RHOB, CEACAM6, TMEM106B, CD44, SLC29A2, SLC7A1, TRAK2, HSF1, MAL2, LRRC8B, PTGES2, VPS41, F2RL1, PTPN6, SLC25A32, YPEL5] |
| **3.85E-04** | GO:1901135 carbohydrate derivative metabolic process | [ALG3, TRIM13, PFKFB2, AK3, ACAT1, ST3GAL1, B4GALNT2, PAICS, ACSF2, RFK, CAD, ELOVL7, CD44, SCD, PFKP] |
| **4.27E-04** | GO:0051179 localization | [F11R, RIMS4, CDH1, SLC39A8, GYG1, SLC16A3, PPFIA3, SLC6A14, SLC5A6, DKC1, MAP1B, SIDT1, SLC27A3, LLGL2, SLC41A2, CYB5R1, KLC2, CRACR2B, ZMAT3, BMP7, CD24, MCU, GRIK3, TUBA1A, MB, HRAS, TIGD5, CAD, TMC6, VPS37B, RHOB, CEACAM6, TMEM106B, CDH13, CD44, SLC29A2, SLC7A1, TRAK2, HSF1, MAL2, LRRC8B, PTGES2, VPS41, F2RL1, PTPN6, SLC25A32, YPEL5] |
| **5.98E-04** | GO:0051188 cofactor biosynthetic process | [RSAD1, PFKFB2, ACAT1, RFK, ELOVL7, SCD, PFKP] |
| **6.05E-04** | GO:0051234 establishment of localization | [F11R, RIMS4, SLC39A8, GYG1, SLC16A3, PPFIA3, SLC6A14, SLC5A6, MAP1B, SIDT1, SLC27A3, LLGL2, SLC41A2, CYB5R1, KLC2, CRACR2B, ZMAT3, CD24, MCU, GRIK3, TUBA1A, MB, HRAS, TIGD5, CAD, TMC6, VPS37B, RHOB, CEACAM6, TMEM106B, CD44, SLC29A2, SLC7A1, TRAK2, HSF1, MAL2, LRRC8B, PTGES2, VPS41, F2RL1, PTPN6, SLC25A32, YPEL5] |
| **7.31E-04** | GO:0006732 coenzyme metabolic process | [PFKFB2, ACAT1, ACSF2, RFK, ELOVL7, SCD, SLC25A32, PFKP] |
| **9.69E-04** | GO:0009058 biosynthetic process | [CYP1B1, PIP4K2C, GYG1, GPT2, RSAD1, ALG3, DKC1, GRWD1, PFKFB2, AK3, ACAT1, CYB5R1, INPP5J, ASL, ST3GAL1, PAICS, NCAPG2, MRPL4, RFK, SREBF1, CAD, ELOVL7, SGMS2, SRM, HSF1, SC5D, TICRR, SCD, PFKP] |
| **9.91E-04** | GO:0098609 cell-cell adhesion | [CDH1, CYP1B1, L1CAM, BMP7, CD24, CEACAM6, CLDN12, CDH13, CD44, PTPN6] |
| **1.04E-03** | GO:0051186 cofactor metabolic process | [GPT2, RSAD1, PFKFB2, ACAT1, ACSF2, RFK, ELOVL7, SCD, SLC25A32, PFKP] |
| **1.08E-03** | GO:1901566 organonitrogen compound biosynthetic process | [GPT2, RSAD1, PFKFB2, AK3, ACAT1, ASL, ST3GAL1, PAICS, RFK, CAD, ELOVL7, SGMS2, SRM, SCD, PFKP] |
| **1.14E-03** | GO:0030036 actin cytoskeleton organization | [F11R, KRT8, LLGL2, SPECC1, TIGD5, RHOB, FGD3, CDC42BPG] |
| **1.15E-03** | GO:0032787 monocarboxylic acid metabolic process | [CYP1B1, SLC16A3, SLC27A3, PFKFB2, ACAT1, ACSF2, ELOVL7, PTGES2, SCD, PFKP] |
| **1.23E-03** | GO:0007162 negative regulation of cell adhesion | [CDH1, CYP1B1, B4GALNT2, CDH13, SEMA3E, ERBB3, PTPN6] |

| | | |
|---|---|---|
| **1.23E-03** | GO:0032956 regulation of actin cytoskeleton organization | [F11R, EPHA1, FMN1, HRAS, TIGD5, RHOB, SEMA3E, F2RL1] |
| **1.23E-03** | GO:1903561 extracellular vesicle | [F11R, CDH1, PIP4K2C, PPFIA3, SLC6A14, KRT8, ACAT1, CYB5R1, ASL, ST3GAL1, PAICS, BPIFB1, HLA-DRB1, TUBA1A, MB, BCAS1, CAD, TMC6, VPS37B, RHOB, CDH13, CD44, MAL2, PTPN6, PFKP] |
| **1.24E-03** | GO:0043230 extracellular organelle | [F11R, CDH1, PIP4K2C, PPFIA3, SLC6A14, KRT8, ACAT1, CYB5R1, ASL, ST3GAL1, PAICS, BPIFB1, HLA-DRB1, TUBA1A, MB, BCAS1, CAD, TMC6, VPS37B, RHOB, CDH13, CD44, MAL2, PTPN6, PFKP] |
| **1.26E-03** | GO:0055086 nucleobase-containing small molecule metabolic process | [PFKFB2, AK3, ACAT1, B4GALNT2, PAICS, ACSF2, RFK, CAD, ELOVL7, SCD, PFKP] |
| **1.28E-03** | GO:0009117 nucleotide metabolic process | [PFKFB2, AK3, ACAT1, PAICS, ACSF2, RFK, CAD, ELOVL7, SCD, PFKP] |
| **1.34E-03** | GO:0044089 positive regulation of cellular component biogenesis | [PIP4K2C, EPHA1, BMP7, FMN1, WDR43, HRAS, TIGD5, MIEN1, HSF1, F2RL1] |
| **1.42E-03** | GO:0006753 nucleoside phosphate metabolic process | [PFKFB2, AK3, ACAT1, PAICS, ACSF2, RFK, CAD, ELOVL7, SCD, PFKP] |
| **1.45E-03** | GO:0007155 cell adhesion | [F11R, CDH1, CYP1B1, EPDR1, EPHA1, L1CAM, BMP7, CD24, RHOB, CEACAM6, CLDN12, CDH13, CD44, PTPN6] |
| **1.46E-03** | GO:1901576 organic substance biosynthetic process | [PIP4K2C, GYG1, GPT2, RSAD1, ALG3, DKC1, GRWD1, PFKFB2, AK3, ACAT1, CYB5R1, INPP5J, ASL, ST3GAL1, PAICS, NCAPG2, MRPL4, RFK, SREBF1, CAD, ELOVL7, SGMS2, SRM, HSF1, SC5D, TICRR, SCD, PFKP] |
| **1.54E-03** | GO:0022610 biological adhesion | [F11R, CDH1, CYP1B1, EPDR1, EPHA1, L1CAM, BMP7, CD24, RHOB, CEACAM6, CLDN12, CDH13, CD44, PTPN6] |
| **1.61E-03** | GO:0016021 integral component of membrane | [F11R, EVA1B, CDH1, SLC39A8, SLC16A3, SLC6A14, ALG3, SLC5A6, TRIM13, SIDT1, EPHA1, SLC27A3, SLC41A2, CYB5R1, TSPAN9, ST3GAL1, B4GALNT2, TMEM120B, TMEM86A, ST14, GLMP, L1CAM, MCU, HLA-DRB1, GRIK3, TMEM181, TMEM47, SREBF1, ELOVL7, TMC6, CLDN12, TMEM106B, PGAM5, CD44, SLC29A2, SEMA3E, SGMS2, SLC7A1, ERBB3, SC5D, MAL2, LRRC8B, TM4SF18, PTGES2, GALNT1, SCD, F2RL1, SLC25A32] |
| **1.62E-03** | GO:0009150 purine ribonucleotide metabolic process | [PFKFB2, AK3, ACAT1, PAICS, ACSF2, ELOVL7, SCD, PFKP] |
| **1.64E-03** | GO:0044425 membrane part | [F11R, RIMS4, EVA1B, CDH1, CYP1B1, SLC39A8, SLC16A3, SLC6A14, KRT8, ALG3, SLC5A6, TRIM13, SIDT1, EPHA1, SLC27A3, SLC41A2, CYB5R1, TSPAN9, ST3GAL1, B4GALNT2, TMEM120B, TMEM86A, ST14, GLMP, L1CAM, CD24, MCU, HLA-DRB1, GRIK3, TMEM181, TMEM47, TUBA1A, SREBF1, ELOVL7, TMC6, VPS37B, RHOB, CEACAM6, CLDN12, TMEM106B, CDH13, PGAM5, CD44, SLC29A2, SEMA3E, SGMS2, SLC7A1, MIEN1, ERBB3, SC5D, MAL2, LRRC8B, TM4SF18, PTGES2, GALNT1, SCD, VPS41, F2RL1, PTPN6, SLC25A32] |
| **1.66E-03** | GO:0032989 cellular component morphogenesis | [TSKU, CDH1, MAP1B, EPHA1, ST14, BMP7, TMEM106B, CDH13, TRAK2] |

| | | |
|---|---|---|
| **1.68E-03** | GO:0040017 positive regulation of locomotion | [EPHA1, BMP7, HRAS, VPS37B, RHOB, CEACAM6, CDH13, SEMA3E, MIEN1, F2RL1] |
| **1.71E-03** | GO:0006629 lipid metabolic process | [CYP1B1, PIP4K2C, ALG3, SLC27A3, ACAT1, CYB5R1, INPP5J, B4GALNT2, TMEM86A, ACSF2, SREBF1, ELOVL7, SGMS2, ERBB3, SC5D, PTGES2, SCD] |
| **2.17E-03** | GO:0040012 regulation of locomotion | [CDH1, CYP1B1, TRIM13, EPHA1, BMP7, HRAS, VPS37B, RHOB, CEACAM6, CDH13, SEMA3E, MIEN1, ERBB3, F2RL1] |
| **2.29E-03** | GO:0006163 purine nucleotide metabolic process | [PFKFB2, AK3, ACAT1, PAICS, ACSF2, ELOVL7, SCD, PFKP] |
| **2.33E-03** | GO:0032970 regulation of actin filament-based process | [F11R, EPHA1, FMN1, HRAS, TIGD5, RHOB, SEMA3E, F2RL1] |
| **2.37E-03** | GO:0070062 extracellular exosome | [F11R, CDH1, PIP4K2C, SLC6A14, KRT8, ACAT1, CYB5R1, ASL, ST3GAL1, PAICS, BPIFB1, HLA-DRB1, TUBA1A, MB, BCAS1, CAD, TMC6, VPS37B, RHOB, CDH13, CD44, MAL2, PTPN6, PFKP] |
| **2.65E-03** | GO:0030029 actin filament-based process | [F11R, KRT8, LLGL2, SPECC1, TIGD5, RHOB, FGD3, CDC42BPG] |
| **2.88E-03** | GO:0030335 positive regulation of cell migration | [EPHA1, BMP7, HRAS, RHOB, CEACAM6, CDH13, SEMA3E, MIEN1, F2RL1] |
| **2.89E-03** | GO:0019637 organophosphate metabolic process | [PIP4K2C, PFKFB2, AK3, ACAT1, INPP5J, PAICS, ACSF2, RFK, CAD, ELOVL7, SGMS2, ERBB3, SCD, PFKP] |
| **3.00E-03** | GO:0044249 cellular biosynthetic process | [CYP1B1, PIP4K2C, GYG1, GPT2, RSAD1, DKC1, GRWD1, PFKFB2, AK3, ACAT1, INPP5J, ASL, ST3GAL1, PAICS, NCAPG2, MRPL4, RFK, SREBF1, CAD, ELOVL7, SGMS2, SRM, HSF1, TICRR, SCD, PFKP] |
| **3.03E-03** | GO:0006928 movement of cell or subcellular component | [F11R, CYP1B1, SLC16A3, MAP1B, EPHA1, KLC2, L1CAM, BMP7, CD24, TIGD5, RHOB, CEACAM6, CDH13, CD44, SEMA3E, TRAK2, F2RL1, PTPN6] |
| **3.20E-03** | GO:0006631 fatty acid metabolic process | [CYP1B1, SLC27A3, ACAT1, ACSF2, ELOVL7, PTGES2, SCD] |
| **3.31E-03** | GO:0045785 positive regulation of cell adhesion | [EPHA1, BMP7, CD24, FMN1, CEACAM6, CDH13, CD44, PTPN6] |
| **3.35E-03** | GO:0044255 cellular lipid metabolic process | [CYP1B1, PIP4K2C, ALG3, SLC27A3, ACAT1, INPP5J, B4GALNT2, TMEM86A, ACSF2, ELOVL7, SGMS2, ERBB3, PTGES2, SCD] |
| **3.35E-03** | GO:0044444 cytoplasmic part | [TES, F11R, CMIP, RIMS4, CDH1, CYP1B1, PIP4K2C, GYG1, PPFIA3, GPT2, RSAD1, KRT8, ALG3, MAP1B, GRWD1, TRIM13, SLC27A3, HOXB7, LLGL2, PFKFB2, AK3, ACAT1, CYB5R1, COMMD8, KLC2, INPP5J, ASL, ST3GAL1, B4GALNT2, TFAP2C, GLMP, PAICS, ACSF2, MCU, GALK2, HLA-DRB1, UBE2M, GRIK3, MRPL4, SPRED2, TUBA1A, NOB1, MB, RFK, HRAS, SREBF1, TIGD5, CAD, ELOVL7, TMC6, VPS37B, RHOB, CEACAM6, TMEM106B, CDH13, PGAM5, CD44, SGMS2, TRAK2, SRM, FGD3, MIEN1, HSF1, SC5D, MAL2, LRRC8B, PTGES2, GALNT1, CDC42BPG, SCD, VPS41, F2RL1, PTPN6, SLC25A32, YPEL5, PFKP] |
| **3.62E-03** | GO:0045296 cadherin binding | [TES, F11R, CDH1, KLC2, PAICS, CDH13, PFKP] |

| | | |
|---|---|---|
| **3.83E-03** | GO:2000147 positive regulation of cell motility | [EPHA1, BMP7, HRAS, RHOB, CEACAM6, CDH13, SEMA3E, MIEN1, F2RL1] |
| **3.90E-03** | GO:0016477 cell migration | [F11R, CYP1B1, SLC16A3, MAP1B, L1CAM, CD24, RHOB, CEACAM6, CDH13, CD44, SEMA3E, F2RL1, PTPN6] |
| **3.95E-03** | GO:0044283 small molecule biosynthetic process | [GPT2, PFKFB2, ACAT1, ASL, RFK, CAD, ELOVL7, SC5D, SCD, PFKP] |
| **4.07E-03** | GO:0006793 phosphorus metabolic process | [PIP4K2C, EPHA1, PFKFB2, AK3, ACAT1, INPP5J, ST3GAL1, B4GALNT2, PAICS, ACSF2, GALK2, RFK, HRAS, CAD, ELOVL7, PGAM5, SGMS2, ERBB3, HSF1, CDC42BPG, SCD, PTPN6, PFKP] |
| **4.15E-03** | GO:0072521 purine-containing compound metabolic process | [PFKFB2, AK3, ACAT1, PAICS, ACSF2, ELOVL7, SCD, PFKP] |
| **4.37E-03** | GO:0046394 carboxylic acid biosynthetic process | [GPT2, PFKFB2, ASL, CAD, ELOVL7, SCD, PFKP] |
| **4.44E-03** | GO:0050730 regulation of peptidyl-tyrosine phosphorylation | [CD24, NCAPG2, CD44, ERBB3, HSF1, PTPN6] |
| **4.44E-03** | GO:0016053 organic acid biosynthetic process | [GPT2, PFKFB2, ASL, CAD, ELOVL7, SCD, PFKP] |
| **4.48E-03** | GO:0051272 positive regulation of cellular component movement | [EPHA1, BMP7, HRAS, RHOB, CEACAM6, CDH13, SEMA3E, MIEN1, F2RL1] |
| **4.80E-03** | GO:0098656 anion transmembrane transport | [SLC16A3, SLC6A14, SLC5A6, SLC7A1, LRRC8B, SLC25A32] |
| **5.20E-03** | GO:0044271 cellular nitrogen compound biosynthetic process | [CYP1B1, RSAD1, DKC1, PFKFB2, AK3, ACAT1, ASL, PAICS, NCAPG2, RFK, SREBF1, CAD, ELOVL7, SGMS2, SRM, HSF1, SCD, PFKP] |
| **5.32E-03** | GO:0030054 cell junction | [TES, F11R, RIMS4, CDH1, KRT8, MAP1B, TSPAN9, L1CAM, GRIK3, FMN1, TMEM47, RHOB, CLDN12, CDH13, CD44, PTPN6] |
| **5.47E-03** | GO:0051493 regulation of cytoskeleton organization | [F11R, MAP1B, EPHA1, FMN1, HRAS, TIGD5, RHOB, SEMA3E, F2RL1] |
| **5.78E-03** | GO:0016310 phosphorylation | [PIP4K2C, EPHA1, PFKFB2, AK3, ST3GAL1, GALK2, RFK, HRAS, CAD, SGMS2, ERBB3, HSF1, CDC42BPG, PTPN6, PFKP] |
| **5.89E-03** | GO:0031090 organelle membrane | [CYP1B1, SLC39A8, SLC16A3, SLC5A6, SLC27A3, CYB5R1, ST3GAL1, B4GALNT2, TMEM120B, GLMP, MCU, HLA-DRB1, MRPL4, SPRED2, HRAS, SREBF1, TMC6, VPS37B, RHOB, CEACAM6, TMEM106B, PGAM5, CD44, SLC29A2, PTGES2, GALNT1, VPS41, SLC25A32] |
| **6.01E-03** | GO:0009987 cellular process | [F11R, RCBTB1, TSKU, RIMS4, CDH1, CYP1B1, SLC39A8, PIP4K2C, GYG1, SLC16A3, PPFIA3, DNAAF3, GPT2, RSAD1, SLC6A14, KRT8, ALG3, OVOL1, DKC1, MAP1B, GRWD1, TRIM13, EPHA1, SLC27A3, HOXB7, LLGL2, PFKFB2, NOLC1, AK3, |

| | | |
|---|---|---|
| | | TRMT2A, CDKN2AIP, ACAT1, CYB5R1, KLC2, ZMAT3, TSPAN9, INPP5J, ASL, ST3GAL1, B4GALNT2, SPECC1, TMEM120B, TMEM86A, TFAP2C, ST14, L1CAM, PAICS, BMP7, ACSF2, BHLHE40, CD24, MCU, GALK2, HLA-DRB1, UBE2M, NCAPG2, GRIK3, FMN1, WDR43, MRPL4, SPRED2, TUBA1A, NOB1, MB, RFK, HRAS, BCAS1, SREBF1, TIGD5, CAD, ELOVL7, TMC6, VPS37B, RHOB, CEACAM6, TMEM106B, CDH13, PGAM5, CD44, SLC29A2, SEMA3E, SGMS2, TRAK2, SRM, FGD3, MIEN1, ERBB3, HSF1, MAL2, LRRC8B, PTGES2, GALNT1, CDC42BPG, TICRR, SCD, VPS41, F2RL1, PTPN6, SLC25A32, YPEL5, PFKP] |
| **6.28E-03** | GO:0032940 secretion by cell | [RIMS4, GYG1, PPFIA3, LLGL2, CYB5R1, TMC6, CEACAM6, CD44, PTGES2, VPS41, F2RL1, PTPN6, YPEL5] |
| **6.32E-03** | GO:0005912 adherens junction | [TES, CDH1, TSPAN9, L1CAM, FMN1, TMEM47, RHOB, CDH13, CD44] |
| **6.36E-03** | GO:0046903 secretion | [RIMS4, GYG1, PPFIA3, LLGL2, CYB5R1, CAD, TMC6, CEACAM6, CD44, PTGES2, VPS41, F2RL1, PTPN6, YPEL5] |
| **6.72E-03** | GO:0031226 intrinsic component of plasma membrane | [SLC39A8, SLC16A3, SLC5A6, EPHA1, TSPAN9, ST14, CD24, GRIK3, TMC6, CD44, SLC29A2, SEMA3E, SGMS2, SLC7A1, MIEN1, ERBB3, LRRC8B, F2RL1] |
| **6.85E-03** | GO:0006887 exocytosis | [RIMS4, GYG1, LLGL2, CYB5R1, TMC6, CEACAM6, CD44, PTGES2, VPS41, PTPN6, YPEL5] |
| **6.89E-03** | GO:0050900 leukocyte migration | [F11R, SLC16A3, L1CAM, CEACAM6, CD44, F2RL1, PTPN6] |
| **6.98E-03** | GO:0120035 regulation of plasma membrane bounded cell projection organization | [MAP1B, L1CAM, BMP7, HRAS, TMEM106B, CD44, SEMA3E, TRAK2, MIEN1, F2RL1] |
| **7.17E-03** | GO:0040011 locomotion | [F11R, CYP1B1, SLC16A3, MAP1B, L1CAM, CD24, HRAS, RHOB, CEACAM6, CDH13, CD44, SEMA3E, F2RL1, PTPN6] |
| **7.18E-03** | GO:0006796 phosphate-containing compound metabolic process | [PIP4K2C, EPHA1, PFKFB2, AK3, ACAT1, INPP5J, ST3GAL1, PAICS, ACSF2, GALK2, RFK, HRAS, CAD, ELOVL7, PGAM5, SGMS2, ERBB3, HSF1, CDC42BPG, SCD, PTPN6, PFKP] |
| **7.18E-03** | GO:0046942 carboxylic acid transport | [SLC16A3, SLC6A14, SLC5A6, SLC27A3, SLC7A1, SLC25A32] |
| **7.18E-03** | GO:0015849 organic acid transport | [SLC16A3, SLC6A14, SLC5A6, SLC27A3, SLC7A1, SLC25A32] |
